# Guiding T lymphopoiesis from pluripotent stem cells by defined transcription factors

**DOI:** 10.1101/660977

**Authors:** Rongqun Guo, Fangxiao Hu, Qitong Weng, Cui Lv, Hongling Wu, Lijuan Liu, Zongcheng Li, Yang Zeng, Zhijie Bai, Mengyun Zhang, Yuting Liu, Xiaofei Liu, Chengxiang Xia, Tongjie Wang, Peiqing Zhou, Kaitao Wang, Yong Dong, Yuxuan Luo, Xiangzhong Zhang, Yuxian Guan, Yang Geng, Juan Du, Yangqiu Li, Yu Lan, Jiekai Chen, Bing Liu, Jinyong Wang

## Abstract

Achievement of immunocompetent and therapeutic T lymphopoiesis from pluripotent stem cells is a central aim in T cell regenerative medicine. To date, preferentially regenerating T lymphopoiesis *in vivo* from pluripotent stem cells (PSC) remains a practical challenge. Here we documented that synergistic and transient expression of Runx1 and Hoxa9 restricted in the time window of endothelial to hematopoietic transition and hematopoietic maturation stages induced *in vitro* from PSC (iR9-PSC) preferentially generated engraftable hematopoietic progenitors capable of homing to thymus and developing into mature T (iT) cells in primary and secondary immunodeficient recipients. Single-cell transcriptome and functional analyses illustrated the cellular trajectory of T lineage induction from PSC, unveiling the T-lineage specification determined at as early as hemogenic endothelial cell stage and identifying the *bona fide* pre-thymic progenitors. The iT cells distributed normally in central and peripheral lymphoid organs and exhibited abundant TCRαβ repertoire. The regenerative T lymphopoiesis rescued the immune-surveillance ability in immunodeficient mice. Furthermore, gene-edited iR9-PSC produced tumor-specific-T cells *in vivo* that effectively eradicated tumor cells. This study provides insight into universal generation of functional and therapeutic T lymphopoiesis from the unlimited and editable PSC source.

## INTRODUCATION

A prominent method to generate T cells is an in vitro system via co-culture of either mouse or human hematopoietic stem/progenitors (HSPC) with stromal cell lines expressing the Notch ligand, such as OP9-DL1/DL4 or 3D-based MS5-hDLL1/4 ^1–3^. Despite the great contribution of this approach to studying T cell development *in vitro*, phenotypic T cells produced by this approach face severe immunocompetency problems *in vivo* after engraftment, due to the inadequate *in vitro* recapitulation of natural thymus microenvironment. Natural mouse Sca1^+^cKit^+^ and human CD34^+^ blood progenitor cells can be induced into CD7^+^ pre-thymic cells *in vitro*, which successfully colonize thymi and mature into immunocompetent T cells *in vivo* ^4, 5^. However, this in vitro plus in vivo two-step approach never succeeded in generating induced T lymphopoiesis when starting from pluripotent stem cells (PSC), as induced T cell progenitors from PSC showed intrinsic thymus-homing defect *in vivo* ^6^. Another prevailing concept for generating functional T lymphopoiesis from PSC is via induction of hematopoietic stem cell (HSC)-like intermediates followed by in vivo multi-lineage hematopoiesis, including T cells ^7–10^. However, generating robust *bona fide* induced-HSC (iHSC) from PSC remains inefficient ^11, 12^ and whether this approach can generate therapeutic tumor-killing T cells is unknown. Recently, Hoxb5 is shown to convert natural B cells into functional T cells *in vivo* ^13^, providing an alternative method to shorten the immune system recovery gap in conventional HSC transplantation. Nonetheless, a solid and universal approach, capable of generating immunocompetent and therapeutic T lymphopoiesis from the unlimited and gene-editable PSC, is still lacking.

Accumulated developmental evidence shows that blood progenitors prior to the occurrence of definitive HSC, also possess T cell lineage differentiation potential ^14–17^. Despite the abundant knowledge of the pivotal transcription factors regulating T cell development from HSC derivatives ^18^, intrinsic determinants of T cell lineage potential in the HSC-independent hematopoietic progenitors at the pre-liver and pre-thymus stages remain elusive. Thus, identifying such crucial T lineage-potential determinants might help to establish a solid protocol for efficiently reconstituting T lymphopoiesis from PSC.

In this study by a unbiased *in vivo* functional screening approach, we identified that the coordinated and transient expression of exogenous *Runx1* and *Hoxa9* at the early time window from endothelial to hematopoietic transition stage to hematopoietic progenitor maturation stage induced in vitro from PSC, produced a type of induced hematopoietic progenitor cells (iHPC) with thymus-homing features, which was engraftable and gave rise to induced T cells (iT cells) with abundant TCRαβ repertoire in immune deficient mice. Physiologically, the iT cells successfully rescued immune surveillance function in immune deficient mice. Therapeutically, these iT cells possessed anti-tumor activities *in vivo* when engineered to carry tumor antigen specific TCR at PSC stage. For the first time, we establish a novel approach of preferentially generating functional and therapeutic T lymphopoiesis *in vivo* from PSC, which technically creates a link between the unlimited and editable PSC source and T cell-based immunotherapy for translational purpose.

## RESULTS

### Reconstitution of T lymphopoiesis *in vivo* from inducible *Runx1-p2a-Hoxa9*-embryonic stem cells

We hypothesized that the lymphogenic potential is determined by intrinsic determinants at putative endothelial precursor cell stage prior to and independent of HSC formation. Therefore, enforced expression of these master determinants might direct lymphoid differentiation from PSC. Since *Runx1* is pivotal for endothelial to hematopoietic transition (EHT) ^19–21^, definitive hematopoietic development ^22–24^ and T cell development ^18^, we started from evaluating the potential effect of *Runx1* in lymphogenic commitment from PSC. To avoid the expression variations introduced by viral delivery systems, we inserted the inducible expression cassette of *Runx1* into the *Rosa26 locus* of embryonic stem cells (*iRunx1*-ESC, C57BL/6 background) by homologous recombination (Supplementary information, Fig. S1a), which resulted in the conditional expression of exogenous *Runx1* in the presence of doxycycline (Supplementary information, Fig. S1b). We used AFT024-(mSCF/mIL3/mIL6/hFlt3L) cell line-cultured supernatants as conditioned medium (CM) for the *in vitro* induction of induced hemogenic endothelial progenitors (iHEC) and subsequent iHPC, as AFT024 CM is beneficial for the generation of induced HPC *in vitro* ^25^. To functionally assess the T lymphopoiesis potential of iHPC, we transplanted the bulk cells containing abundant iHPC (referred as iHPC thereafter) into irradiated (2.25 Gy) B-NDG recipients (iHPC recipients) and used the occurrence of CD3^+^ cells in peripheral blood (PB) as a positive readout of induced T lymphopoiesis *in vivo* (Fig. 1a). Based on a modified protocol for HEC induction from PSC ^26^, we successfully generated iHEC and hematopoietic progenitor derivatives (Supplementary information, Figs. S1c-e). However, the *iRunx1*-ESC derivatives eventually failed to generate T cells on the conditions of either *in vitro* OP9-DL1 co-culture system (Supplementary information, Fig. S1f) or *in vivo* transplantation setting (Supplementary information, Fig. S1g). We speculated that the other transcription factors essential for T lineage generation might be absent in the *iRunx1*-ESC derivatives. To identify these absent factors, we sorted the single iHEC from *iRunx1*-ESC and performed single-cell RNA-Seq. In comparison with E11 T1-pre-HSC (CD31^+^CD41^low^CD45^−^c-kit^+^CD201^high^), we identified eight hematopoietic-essential transcription factors, *Hoxa5* ^8^, *Hoxa7* ^27^, *Hoxa9* ^28^, *Hoxa10* ^29^, *Hlf* ^30^, *Ikzf1* ^31^, *Nkx2-3* ^32^, and *Setbp1* ^33^, which were barely expressed in *iRunx1*-ES-derived iHEC but abundantly expressed in E11 T1-pre-HSC (Fig. 1b). Consistent with the previous reports that human PSC-derived HEC lacks expression of HOXA family ^8, 34^. We further used an “*iRunx1*+*Xi*” tandem-factor-knock-in strategy to perform unbiased screening of the potential combinatory effects of these factors with Runx1 in lymphoid lineage induction. Following the same induction protocol, we identified that the inducible expression of exogenous *Runx1* and *Hoxa9* from day 6 to day 11 during the induction program led to the production of robust iHEC phenotypically resembling embryonic pre-HSC (CD31^+^CD41^low^CD45^−^c-kit^+^CD201^high^) (Fig. 1c) ^35^. Notably, CD201^+/high^ expression can enrich hemogenic precursors with both definitive HPC and HSC potential from as early as E9.5 embryos ^36^. After co-culture of these iHEC with OP9-DL1 feeder line (GFP^+^) in the presence of CM and doxycycline (1 μg/ml), robust iHPC occurred at day 21, including phenotypic pre-thymic progenitors (Lin^−^c-kit^+^CD127^+^/CD135^+^) ^18^ (Fig. 1d), and CD11b^+^/Gr1^+^ myeloid cells, but no CD3^+^ T cells (Supplementary information, Fig. S1h). To further assess the engraftment potential of these iHPC, we transplanted 0.5-1 million *iR9*-ESC-derived iHPC (day-21) into irradiated (2.25 Gy) B-NDG mice (8-week-old, CD45.1 strain) in the absence of doxycycline. Four weeks after transplantation, we observed donor-derived CD45.2^+^ CD3^+^ T cells, but no CD45.2^+^CD19^+^ B cells and no CD45.2^+^CD11b^+^ myeloid cells, in the PB of B-NDG mice transplanted with the iHPC (Fig. 1e). Five independent experiments indicated that the *iR9*-ESC-derived iHPC gave rise to CD3^+^ iT cells in over 80% B-NDG recipients (iT-B-NDG mice, 32/40) (Fig. 1f; Supplementary information, Fig. S1i). In addition, the day-17 iHPC also reconstituted T lymphopoiesis in B-NDG recipients (Supplementary information, Figs. S2a-d). Thus, we established a novel approach of preferentially generating iT cells from gene-edited PSC by defined transcription factor *Runx1* and *Hoxa9*.

**Fig. 1.**
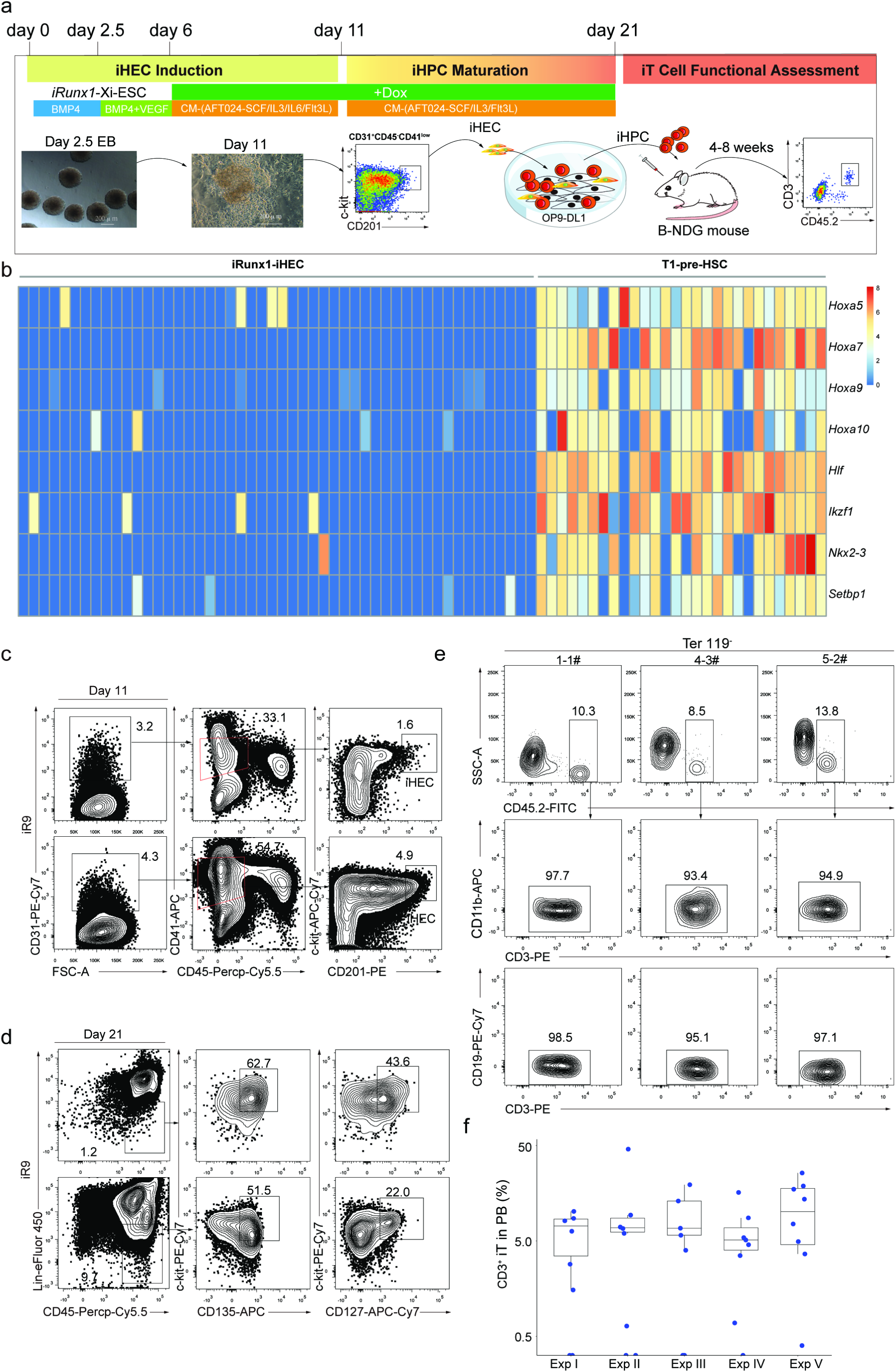
T cell regeneration *in vivo* from *iRunx1-p2a-Hoxa9*-edited embryonic stem cells. **a** The strategy of stepwise T lineage induction by defined transcription factors. iRunx1-ESC, and iRunx1-Xi-ESC lines (C57BL/6 background, CD45.2 strain) were used for T cell lineage induction. Xi means one of the eight transcription factors *Hoxa5*, *Hoxa7*, *Hoxa9*, *Hoxa10*, *Hlf*, *Ikzf1*, *Nkx2-3*, *Setbp1*. **b** Heatmaps of the eight transcription factors abundantly expressed in embryonic pre-HSC but rarely expressed in iRunx1-ES derived iHEC. The expression value (TPM) of each gene was converted by log2 and illustrated by pheatmap (R package). One column represents one cell repeat. (iRunx1-iHEC, n=50 single cells, T1-pre-HSC, n=28 single cells). **c** Sorting gates of iHEC population at day 11 derived from *iRunx1-Hoxa9*-ES line (*iR9*-ESC). Two representative plots from five independent experiments are shown. **d** Immuno-phenotypes of pre-thymic progenitors in induced hematopoietic progenitor cells from iHEC after ten-day maturation. Two representative plots from five independent experiments. Lin was defined as CD2^−^CD3^−^CD4^−^CD8^−^CD11b^−^Gr1^−^Ter119^−^CD19^−^NK1.1^−^TCRγδ^−^. Pre-thymic progenitors were defined as Lin^−^c-kit^+^CD127^+^/CD135^+^. **e** Pluripotent stem cell-derived T cells in PB of B-NDG mice were analyzed by flow cytometry 4 weeks after transplantation. One million iHEC-derived hematopoietic cells were transplanted into individual B-NDG mice (CD45.1^+^) irradiated by X-ray (2.25 Gy). Three representative mice from five independent experiments were analyzed. **f** Summary of pluripotent stem cell-derived T cells in PB of individual B-NDG mice from five independent experiments. Forty B-NDG mice transplanted with ESC-derived iHPC were analyzed. The box plot shows the percentage of the CD3^+^ iT cells in PB, the percentage values were illustrated by ggplot2 (R package). A base-10 logarithmic scale was used for the Y-axis. One point represents one mouse.

### The iT cells show features of multi-organ distributions and abundant TCR diversity

We further analyzed the tissue distributions and immunophenotypes of the regenerated T lymphocytes in iT-B-NDG mice. Mature CD4SP and CD8SP iT cells were detected in the spleen, lymph node and PB of iT-B-NDG mice, the majority of which were TCRβ positive (Fig. 2a). In addition, γδ iT cells were also detected in gut and lung tissues of iT-B-NDG mice (Supplementary information, Fig. S3a). Induced NK cells (iNK, CD45.2^+^NK1.1^+^CD3^−^) were also detected in the spleen and bone marrow of iT-B-NDG mice (Supplementary information, Fig. S3b). The thymus of iT-B-NDG mice also contained induced CD4SP (iCD4SP), induced double positive (iDP, CD45.2^+^CD4^+^CD8^+^), induced CD8SP (iCD8SP), and induced double negative (iDN, CD45.2^+^Lin^−^CD4^−^CD8^−^) cells when examined at week-4 and week-5 after transplantation of iHPC. Interestingly, the majority of the iDN cells were at iDN1 (CD45.2^+^Lin^−^CD4^−^CD8^−^CD44^+^CD25^−^) phase at week-4, and at iDN2 (CD45.2^+^Lin^−^CD4^−^CD8^−^CD44^+^CD25^+^)/iDN3 (CD45.2^+^Lin^−^CD4^−^CD8^−^CD44^−^CD25^+^) phases at week-5 (Fig. 2b). Besides the iT cells and induced NK1.1^+^CD3^−^ NK (iNK) cells detected in bone marrow, we also observed *iR9*-ES-derived Lin^−^Sca1^+^cKit^+^ (iLSK) progenitor cells (Fig. 2c). To assess whether the iLSK cells can contribute to T lymphopoiesis, we sorted this population from primary iHPC recipients (week-6) and performed secondary transplantation. Six weeks after transplantation, iT cells appeared in PB, BM, and SP of the B-NDG recipients (Fig. 2d). Of note, despite *iR9*-ES-derived myeloid lineage cells were barely detected *in vivo*, the iLSK cells indeed gave rise to very limited myeloid colonies in CFU assay (data not shown). To further characterize the iT cells at transcriptome level, we sorted 1,000 cell aliquots of the CD4SP iT cells and CD8SP iT cells from the spleens of iT-B-NDG mice for RNA-Seq analysis. Our data indicated that the CD4SP iT cells resembled natural CD4SP T cells, and the CD8SP iT cells resembled natural CD8SP T cells, both of which expressed surface marker-encoding genes *Cd2*, *Fas*, *Cd3e*, *Cxcr3*, *Cd28*, *Cd27*, *Cd7*, *Cd5*, and *Il7r* (Fig. 2e). Of note, the CD4SP iT cells, but not CD8SP iT cells, expressed the *ThPOK* (T helper inducing POK factor, also known as *Zbtb7b*), a master regulator in regulating CD4 vs. CD8 T cell lineage commitment ^37^. In addition, the iT cells also expressed T cell identity genes and key regulators *Tcf7* ^38^, *Tox* ^39^, *Lck* ^40^, *Gata3* ^41^, *Bcl11b* ^42^, *Ikzf2* ^43^, and *Rora* ^44^ (Fig. 2f). In comparison with natural T cell counterparts, the iT cells also showed features of discrepantly expressed genes (a difference in expression of over two-fold; adjusted P value < 0.05 (DESeq2 R package)) (Supplementary information, Table S1), including weaker expression of *Tcf7*. Genomic PCR sequencing using primer pairs flanking the *Runx1-p2a-Hoxa9* element further confirmed that the reconstituted iT cells *in vivo* were of *iR9*-PSC origin, which carried the inserted *Runx1-p2a-Hoxa9* (Supplementary information, Fig. S3c). To further assess the diversities of the TCRαβ clonotypes of the iT cells, we performed TCR deep sequencing using the sorted naïve CD4SP (CD45.2^+^CD4^+^CD62L^+^CD44^−^) and CD8SP iT cells (CD45.2^+^CD8^+^CD62L^+^CD44^−^) from the spleens and thymi of iT-B-NDG mice at week-6 after transplantation of iHPC. The aliquots of 15,000 sorted naïve CD4SP and CD8SP iT cells were used as cell inputs for TCRαβ sequencing at the transcription level. TCRαβ clonotype profiling using MiXCR ^45^ captured abundant diversities of TCRαβ sequences among the sorted naïve iT cells isolated from the thymi (Figs. 2g, h) and spleens (Figs. 2i, j) of the iT-B-NDG mice. Collectively, these data indicate that the *iR9*-ESC-derived iHPC reconstitute T lymphopoiesis *in vivo* resembling natural T cell development.

**Fig. 2.**
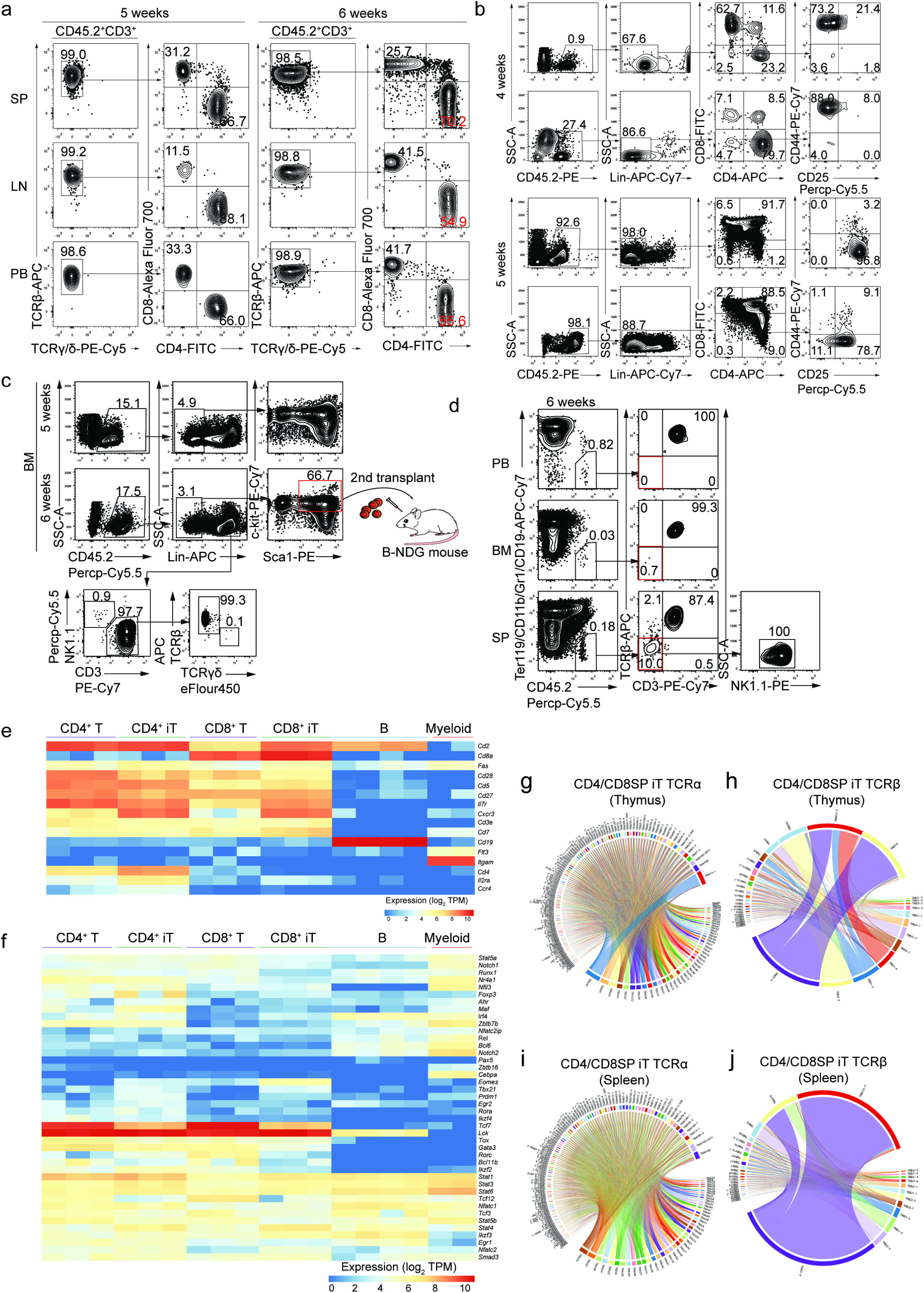
Tissue distributions, transcriptome characterization, and TCRα/β diversities of ESC-derived T Cells. **a** Flow cytometry analysis of mature iT cells in spleen (SP), lymph node (LN), and peripheral blood (PB) of B-NDG mice transplanted with ESC-derived hematopoietic cells. Each B-NDG mouse was transplanted with one million iHPC collected on day 21. Representative mouse was sacrificed and analyzed at 5 and 6 weeks after transplantation. Data from two representative mice are shown. **b** Flow cytometry analysis of iDN cells in the thymus of B-NDG mice transplanted with ESC-derived hematopoietic cells. Each B-NDG mouse was transplanted with one million iHPC at day 21. Representative mouse was sacrificed and analyzed at 4 and 5 weeks after transplantation. Data from four representative mice of two independent experiments are shown. Lin was defined as Ter119^−^CD11b^−^Gr1^−^CD19^−^B220^−^NK1.1^−^TCRγδ^−^. **c** Flow cytometry analysis of iHPC in bone marrow (BM) transplanted with iHPC. Each B-NDG mouse was transplanted with one million iHPC collected at day 10 in the presence of OP9-DL1 feeder cells. Representative mouse was sacrificed and analysed 5 weeks and 6 weeks after transplantation. The BM-derived iHPC (CD45.2^+^Lin^−^c-kit^mid^Sca1^+^) were sorted for 2nd transplantation. Data from two mice are shown. **d** Flow cytometry analysis of iT and iNK in PB, spleen (SP) and bone marrow (BM) 6 weeks after 2nd transplantation. 500 LSK cells from primary iT mice were used as input for secondary transplantation. The secondary recipients were sacrificed and analyzed 6 weeks after transplantation. Data from one mouse are shown. **e** Characterization of surface markers on CD4SP and CD8SP iT cells. CD4SP and CD8SP iT cells were sorted from the spleens of B-NDG mice transplanted with ESC derived hematopoietic cells at week-5. One biological replicate per column. Myeloid cells (n = 2 sample repeats): Ter119^−^CD3^−^CD19^−^CD11b^+^; B cells (n = 4 sample repeats): Ter119^−^CD11b^−^CD3^−^CD19^+^; CD4^+^ cells (n = 3 sample repeats): Ter119^−^CD19^−^CD11b^−^CD4^+^; CD8^+^ cells (n = 3 sample repeats): Ter119^−^CD19^−^ CD11b^−^CD8^+^ iCD4^+^ cells (n = 3 sample repeats): CD45.2^+^Ter119^−^CD19^−^CD11b^−^ CD4^+^; iCD8^+^ cells (n = 3 sample repeats): CD45.2^+^Ter119^−^CD19^−^CD11b^−^CD8^+^. **f** Characterization of transcription factors in CD4SP and CD8SP iT cells. **g** Chord diagram of TCRα diversity in thymus iT cells. **h** Chord diagram of TCRβ diversity in thymus iT cells. **i** Chord diagram of TCRα diversity in thymus iT cells. **j** Chord diagram of TCRβ diversity in spleen iT cells. Aliquots of sorted 15,000 naïve CD4SP and CD8SP iT cells from either thymus or spleen of iT-B-NDG mice were used as cell inputs for TCRαβ sequencing.

### Single iHEC efficiently give rise to iT cells both *in vitro* and *in vivo*

To further investigate the efficiency of iHEC differentiating into iT cells, we sorted single iHEC into individual wells (24 well-plates) pre-seeded with OP9-DL1 feeder cells (Fig. 3a). After ten-day co-culture, over 15 percent individual iHEC formed blood colonies (76/384 wells) (Fig. 3b), which contained abundant pre-thymic progenitors (Lin^−^c-kit^+^CD127^+^/CD135^+^) (Supplementary information, Fig. S4). After co-culture with OP9-DL1 feeder line in the presence of hFlt3L and hIL7, these iHEC-formed blood colonies (30/30) further differentiated into CD3^+^ iT cells *in vitro* (Fig. 3b), including a major population of TCRγδ iT cells, and a small proportion of CD8^+^ TCRβ iT cells (Fig. 3c). To assess the T lymphopoiesis potential of these single-iHEC-derived iHPC, we further collected the iHPC from each colony at day 21 and transplanted them into individual B-NDG mice. Four weeks after transplantation, CD11b^−^CD19^−^CD3^+^ iT cells were detected in approximately 28% (7/25) B-NDG mice transplanted with the cell derivatives from individual iHEC-formed clones (Figs. 3b, d). Collectively, the *iR9*-ESC-derived iHEC robustly gave rise to T cells at the single cell level.

**Fig. 3.**
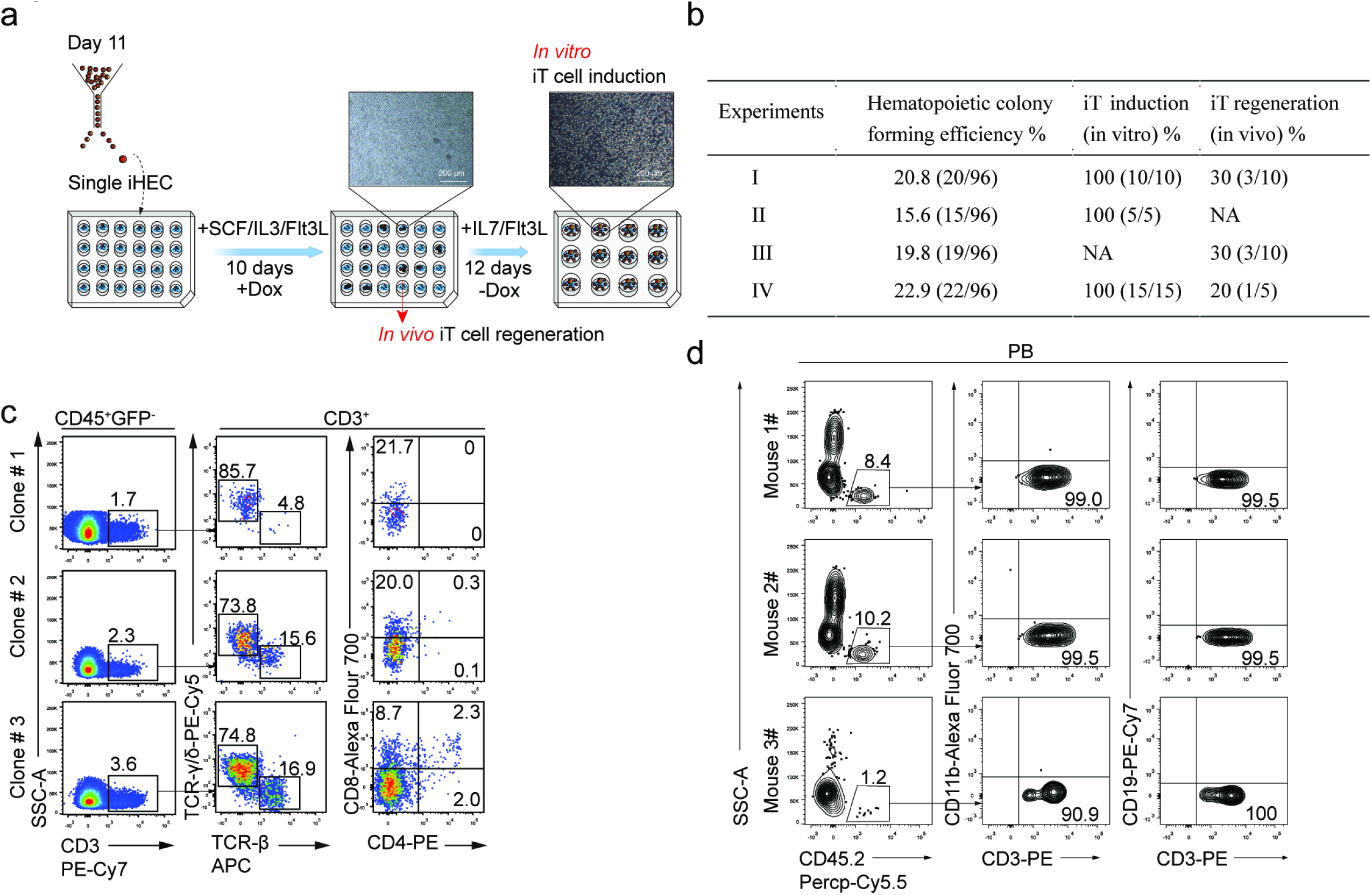
Assessment of T potential of single iHEC from *iR9*-ESC. **a** The strategy of T cell induction from *iR9*-ESC-derived single iHEC. Single iHEC were sorted into individual wells (24-well plates) pre-seeded with OP9-DL1 feeder cells (10000 cells/well) 12 hours prior maturation in EM medium with doxycycline (1 μg/ml). Doxycycline was sustained for 10 days during the maturation step. After maturation, the bulk blood cells were assessed for T lineage generation potential. For in vivo T cell regeneration, the single iHEC-derived bulk hematopoietic cells (day 10) were transplanted into individual B-NDG recipients. For in vitro T cell induction, the medium was changed to T cell induction medium (TIM, α-MEM, 20% DFBS, and 1% GlutaMAX) supplemented with 2% conditioned medium derived from supernatants of AFT024-hFlt3L and AFT024-hIL7 cell culture for sustaining 12 days. **b** Single iHEC efficiently gave rise to T cells. Three hundred and eighty-four single-iHEC at Day 11 were sorted into individual wells (24 well plates). Thirty single-iHEC-formed blood colonies were induced for T cell generation *in vitro*. Cell collections of Twenty-five single-iHEC-formed blood colonies were transplanted into 25 individual B-NDG mice for the assessment of T lymphopoiesis *in vivo*. **c** Flow cytometry analysis of induced T cells from in vitro induction of single iHEC. iT cells from single iHEC culture product (day 22) were analyzed. Plots of iT cells induced from one representative colony are shown. **d** Single iHEC-derived hematopoietic cells gave rise to mature iT cells in PB of B-NDG recipient mice 4 weeks after transplantation. Plots of one representative mouse are shown.

### Cellular trajectory from iHEC to pre-thymic progenitors

To characterize the single iHEC at transcriptome level, we performed single-cell RNA-Seq using the sorted iHEC and compared them with natural single E11 endothelial cells (EC), Type-I pre-HSC, Type-II pre-HSC, E12 HSC, E14 HSC, and adult HSC described previously ^35^. Principle component analysis indicated that the iHEC localized between embryonic EC and pre-HSC (Figs. 4a, b). A large proportion of iHEC expressed artery or vein-related genes, suggestive of their EC-like nature (Fig. 4c). Most iHEC expressed endothelial surface marker-encoding genes *Cdh5* (coding VE-Cadherin, 70/70) and *Esam* (57/70), which were continuously expressed from embryonic EC to pre-HSC at a relatively high level. On the other hand, partial iHEC expressed *Procr* (coding CD201, 32/70), *Cd47* (33/70) and *Cd63* (44/70), which were upregulated from EC to pre-HSC (Fig. 4d). The expression of transcription factors related to endothelial and hematopoietic development further revealed that the iHEC shared a similar feature with embryonic EC and pre-HSC. Majority of the iHEC expressed *Fli1* (66/70), *Erg* (42/70), *Lmo2* (49/70), *Mycn* (65/70), and *Sox7* (38/70). Specifically, a small proportion of iHEC expressed *Bcl11a* (11/70) and *Hoxb5* (24/70). All these transcription factors are pivotal for lymphoid lineage development (Fig. 4e). Thus, the molecular features of the iHEC show similarities with embryonic EC and pre-HSC.

**Fig. 4.**
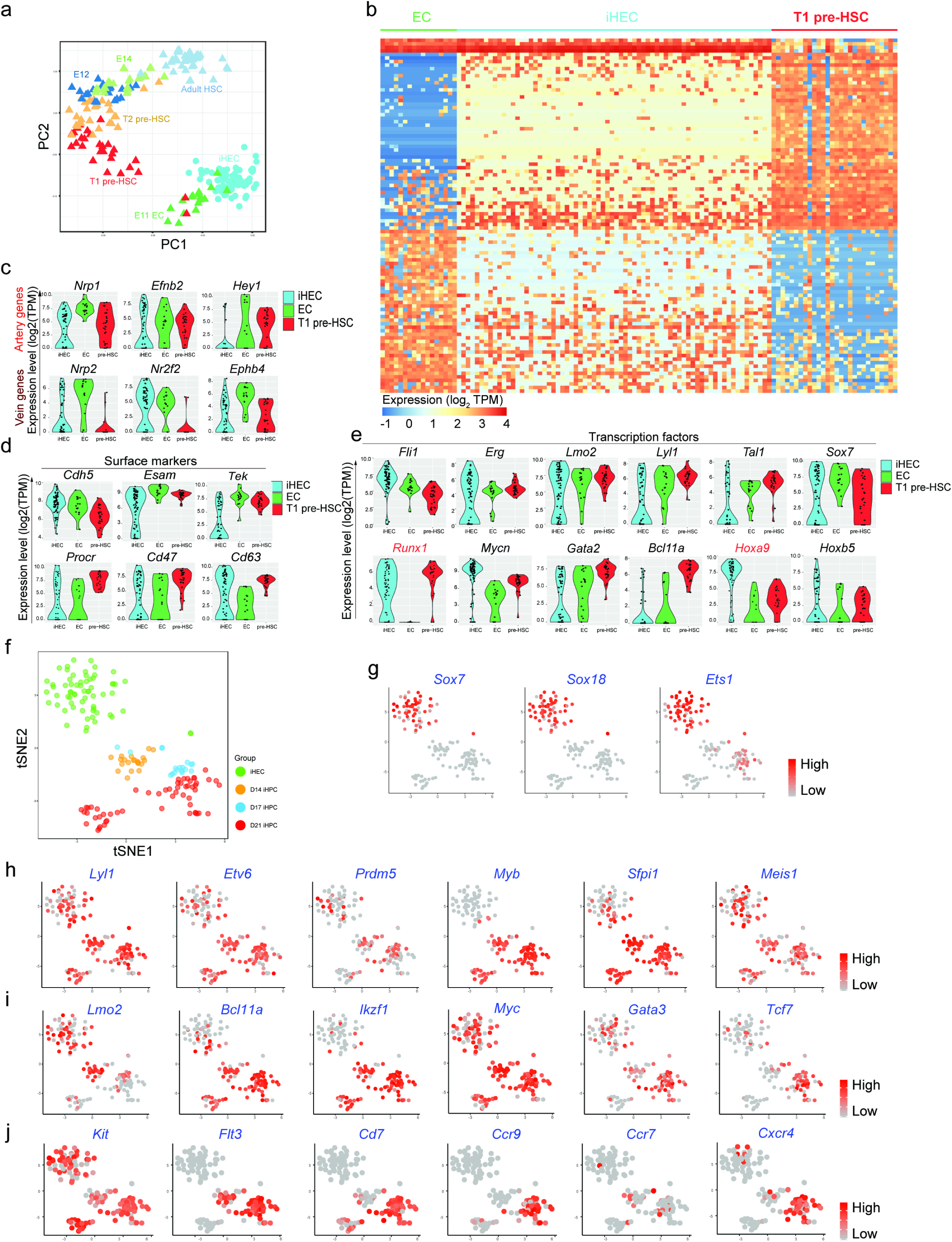
Single-cell transcriptomic characterization of iHEC and iHPC. **a** Principal component analysis (PCA) of iHEC and developmental E11 AGM-derived EC, T1 pre-HSC, T2 pre-HSC, E12 HSC, E14 HSC, and adult HSC. The TPM values of iHEC (n = 70), natural E11 AGM-derived EC (n = 17), T1 pre-HSC (n = 28), T2 pre-HSC (n = 32), E12 HSC (n = 21), E14 HSC (n = 32) and adult HSC (n = 47) single-cell RNA-Seq data were calculated with Stringtie package. **b** The expression of the top 100 genes contributing most to PC2 (50 genes for each direction). The expression value (TPM) of each gene was converted by log2 and illustrated by pheatmap (R package). One column represents one cell repeat. **c** Violin plots show the expression profile of selected artery (A) and vein (V) related genes (A: *Nrp1*, *Efnb2*, and *Hey1*; V: *Nrp2*, *Nr2f2*, and *Ephb4*) in single iHEC. The expression value (TPM) of each gene was converted by log2 and illustrated by ggplot2 (R package). One point represents one cell. **d** Violin plots show the expression profile of selected surface markers (*Cdh5*, *Esam*, *Tek*, *Procr*, *Cd47*, and *Cd63*) in single iHEC. The expression value (TPM) of each gene was converted by log2 and illustrated by ggplot2 (R package). One point represents one cell. **e** Violin plots show the expression profile of selected transcription factors (*Fli1*, *Erg1*, *Lmo2*, *Lyl1*, *Tal1*, *Sox7*, *Runx1*, *Mycn*, *Gata2*, *Bcl11a*, *Hoxa9*, and *Hoxb5*) related to hematopoietic development in single iHEC. The expression value (TPM) of each gene was converted by log2 and illustrated by ggplot2 (R package). One point represents one cell. **f** Two-dimensional tSNE analysis of iHEC and iHPC single-cell RNA-Seq. For single-cell RNA-Seq, the iHEC were collected on day 11, and the iHPC were collected at Day14, 17 and 21. Each dot represents one cell. The TPM values of iHEC (n = 65), iHPC at Day14 (n = 21), Day17 (n = 18) and Day21 (n = 56) from single-cell RNA-Seq data were calculated with Stringtie package. Cell types were defined as: iHEC CD31^+^CD41^low^CD45^−^c-kit^+^CD201^high^; Day14 and Day17 iHPC, CD45^+^Lin (Ter119/Gr1/F4-80/CD2/CD3/CD4/CD8/CD19/FcεRIα)^−^; Day21 iHPC Ter119^−^CD45^+^c-kit^+^ CD127^+^. **g** tSNE analysis of the expression pattern of selected endothelia-related transcription factors (*Sox7*, *Sox18*, and *Ets1*) in iHEC and iHPC. **h** tSNE analysis of the expression pattern of selected hematopoietic-related transcription factors (*Lyl1*, *Etv6*, *Prdm5*, *Myb*, *Sfpi1*, and *Meis1*) in iHEC and iHPC. **i** tSNE analysis of the expression pattern of selected T cell development-related transcription factors (*Lmo2*, *Bcl11a*, *Ikzf1*, *Myc*, *Gata3*, and *Tcf7*) in iHEC and iHPC at Day14, Day17, and Day21. **j** tSNE analysis of the expression pattern of selected lymphopoiesis-related surface protein-coding genes (*Kit*, *Flt3*, *Cd7*, *Ccr9*, *Ccr7*, and *Cxcr4*) in iHEC and iHPC at Day14, Day17, and Day21.

To further characterize the iHPC during the hematopoietic maturation process, we sorted the single iHPC from day-14, day-17, day-21 cell products derived from *iR9*-ES and performed single-cell RNA-Seq. To visualize the time course data of iHPC, we performed t-distributed stochastic neighbor embedding (tSNE, the genes with expression value TPM >1 in more than 30 samples were selected) analysis and illustrated that the day-14-iHPC formed a unique population distinct from day-11-iHEC and the major population of day-17 iHPC. However, the day-17 iHPC and day-21 iHPC already merged (Fig. 4f). In addition, the day-21 iHPC formed a new subpopulation labeled with relatively abundant *Gata2* expression (Supplementary information, Fig. S5a), indicating the heterogeneity of the iHPC. The endothelia-related transcription factors, such as Sox7 and Sox18, were abundantly expressed in day-11 iHEC, however, were immediately silenced in day-14 iHPC (Fig. 4g). The *Ets1* gene, involving embryonic endothelial and lymphoid development ^46^, was shut down in day-14 iHPC but turned on again in day-17 iHPC (Fig. 4g). The transcription factors involving hematopoietic development, such as *Lyl1* ^47^, *Etv6* ^48^, *Prdm5* ^9^, *Myb* ^49^, *Sfpi1* ^50–52^, and *Meis1* ^53^, were widely expressed among day-14, 17, and 21 iHPC populations. (Fig. 4h). Further, the transcription factors related to lymphoid development, including *Lmo2* ^54^, *Bcl11a* ^55^, *Ikzf1* ^56^, *Myc* ^18, 57^, *Gata3* ^58^, and *Tcf7* ^38^, were also expressed in iHPC (Fig. 4i; Supplementary information, Fig. S5b). Of note, day-17 and day-21 iHPC showed abundant expression of *Tcf7* (Fig. 4i). Given the thymus-homing problem of the PSC-derived HPC reported by others ^6^, we observed that the day-21-iHPC derived from *iR9*-ES abundantly expressed surface marker-encoding gene *Kit* ^18^, *Flt3* ^18^, *Cd7* ^4, 5^, *Ccr9* ^59, 60^, and *Cxcr4* ^61, 62^, which is a feature of natural pre-thymic progenitors possessing thymus-homing ability (Fig. 4j). However, the subpopulation with relatively abundant Gata2 expression on Day21 lacks the thymus-homing feature genes, indicating that these cells unlikely contributed to the regenerated iT lymphopoiesis. Collectively, the *iR9*-ES-derived iHPC show hematopoietic or lymphopoietic features at transcriptome level and the day-21 iHPC contain robust pre-thymic progenitor-like cells.

### The iT cells reject allogeneic skin and form memory response *in vivo*

To investigate the function of iT cells derived from *iR9*-ESC (C57BL/6 background) *in vivo*, we transferred the iT cells (5 million equivalents of iT cells per *Rag1^−/−^*) isolated from iT-B-NDG spleen into *Rag1*^−/−^ mice (iT-*Rag1*^−/−^ mice). Four days after the adoptive iT cell transfer, we transplanted allogeneic skin from BALB/c mice into the iT-*Rag1*^−/−^ mice. The allogeneic skin grafts were rapidly rejected by iT-*Rag1*^−/−^ mice at around day 9 after transplantation, as indicated by bulged, ulcerative and necrotic lesions at the graft sites (Fig. 5a). Besides the mature iT cells (CD4SP, CD8SP) in the PB of iT-*Rag1*^−/−^ mice (Fig. 5b), activated CD4SP and CD8SP iT cells (CD44^high^CD69^+^) were also detected in the rejected allogeneic skin tissues (Fig. 5c). The iT-*Rag1^−/−^* mice still showed the existence of iT cells in PB thirty days after the primary allogeneic rejection, and again rejected the secondary allogeneic skin grafts (Supplementary information, Fig. S6). Flow cytometry indicated that IL17^+^ and IFNγ^+^ CD4^+^ iT cells, and IFNγ^+^ CD8^+^ iT cells existed in the primary- and secondary-rejected skin grafts (Fig. 5d). Collectively, these results indicated that the adoptively transferred iT cells in *Rag1*^−/−^ mice mediated rejection of allogeneic skin grafts and sustained immunological memory, suggestive of a typical adaptive immune response.

**Fig. 5.**
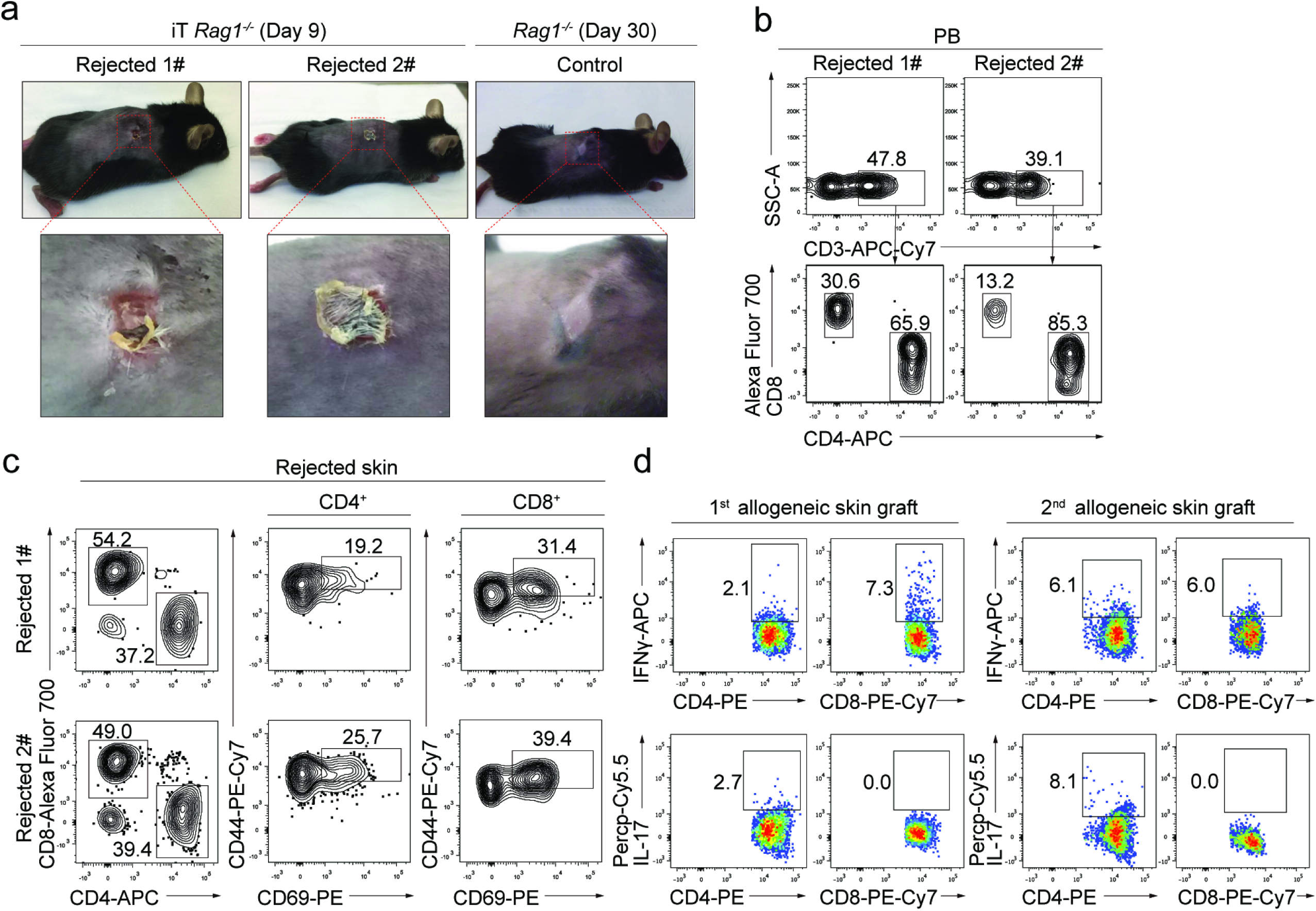
iT cells reject allogeneic skin in adoptively transferred *Rag1^−/−^* mice. **a** The images of allogeneic skin grafts. Representative images of rejected allogeneic skin tissues on ESC-iT-*Rag1^−/−^* (day 9) mice (n = 2) and grafted skin tissue on control *Rag1^−/−^* mice (day 30) were shown. **b** Flow cytometry analysis of the ESC-iT cells in peripheral blood (PB) of adoptively ESC-iT transferred *Rag1^−/−^* recipients nine days after the allogeneic skin grafted. Plots of two representative mice are shown. **c** Flow cytometry analysis of the activation status of the ESC-iT cells in the rejected allogeneic skin tissues. The rejected allogeneic skin tissues were from the adoptively ESC-iT transferred *Rag1^−/−^* recipients nine days after the allogeneic skin grafted. The activated ESC-iT cells were defined as CD4^+^/CD8^+^ CD44^high^ CD69^+^. Rejected skin tissues from two representative ESC-iT transferred *Rag1^−/−^* mice were analyzed. **d** Flow cytometry analysis of the intracellular cytokine IFNγ and IL-17 secreted by the CD4^+^ or CD8^+^ ESC-iT cells in rejected allogeneic skin tissues. 1^st^ allogeneic skin grafts were analyzed at day 9 and 2^nd^ allogeneic skin grafts were analyzed at day 6 after skin transplantation. Data from primary and secondary rejected skin tissues from one representative ESC-iT cells transferred *Rag1^−/−^* mouse are shown.

### The iT cells derived from TCR-edited iPSC eradicate tumor cells *in vivo*

Regarding the advantages of unlimited cell source and gene-editing advantage of iPSC, we introduced tumor antigen-specific TCR (MHC-I restricted OVA TCR, OT1) into *iR9*-iPSC and further assessed the anti-tumor activity of the derived OT1 iT cells. We reprogrammed mouse MEF (C57BL/6 background, CD45.2 strain) into iPSC using retroviruses carrying *Oct4*/*Klf4*/*Sox2*. Two cassettes of *rtTA-TRE-Runx1-p2a-Hoxa9-HygroR* and *CAG-OT1-TCR-IRES-GFP-PuroR* were inserted into the loci of *Rosa26* and *Hipp11* of iPSC (*OT1*-*iR9*-iPSC), respectively (Fig. 6a). Intracellular staining indicated that the OT1-TCR were expressed in the *OT1*-*iR9*-iPSC (Fig. 6b). The *OT1*-*iR9*-iPSC were further induced into OT1-iHEC (Fig. 6c) and OT1-iHPC (Fig. 6d). We transplanted the OT1-iHPC (three million per mouse) into irradiated (4.5 Gy) *Rag1^−/−^* mice (OT1-iHPC recipients) to reconstitute OT1-iT lymphopoiesis. Six weeks after transplantation, the OT1-iHPC recipients showed GFP^+^CD8^+^ iT cells expressing OT1 TCRαβ in PB (Fig. 6e). We then engrafted E.G7-OVA tumor cells into the groin of the *Rag1^−/−^* or OT1-iT reconstituted *Rag1^−/−^* mice (OT1-iT-*Rag1^−/−^* mice) by subcutaneous injection (0.2 million/mouse). Tumor growth kinetics demonstrated that the E.G7-OVA tumors were dramatically inhibited in the OT1-iT-*Rag1^−/−^* mice in comparison with the control *Rag1^−/−^* mice (Fig. 6f). We sacrificed the OT1-iT-*Rag1^−/−^* mice for the distribution analysis of the iT cells in tumors and lymphoid organs 19 days after the tumor cell transplantation. Flow cytometry analysis demonstrated that the E.G7-OVA tumors in the OT1-iT-*Rag1^−/−^* mice were infiltrated with CD8^+^ OT1-iT cells, which contained effector (CD44^+^ CD62L^−^) and memory (CD44^+^ CD62L^+^) iT cells, and IFNγ-secreting iT cells (Fig. 6g). We also observed abundant CD8^+^ iT cells carrying OT1 TCRαβ in the bone marrow, lymph node, and spleen of these mice (Supplementary information, Fig. S7). Collectively, these data indicate that the iT cells derived from TCR-engineered iPSC show anti-tumor activity in a solid tumor model.

**Fig. 6.**
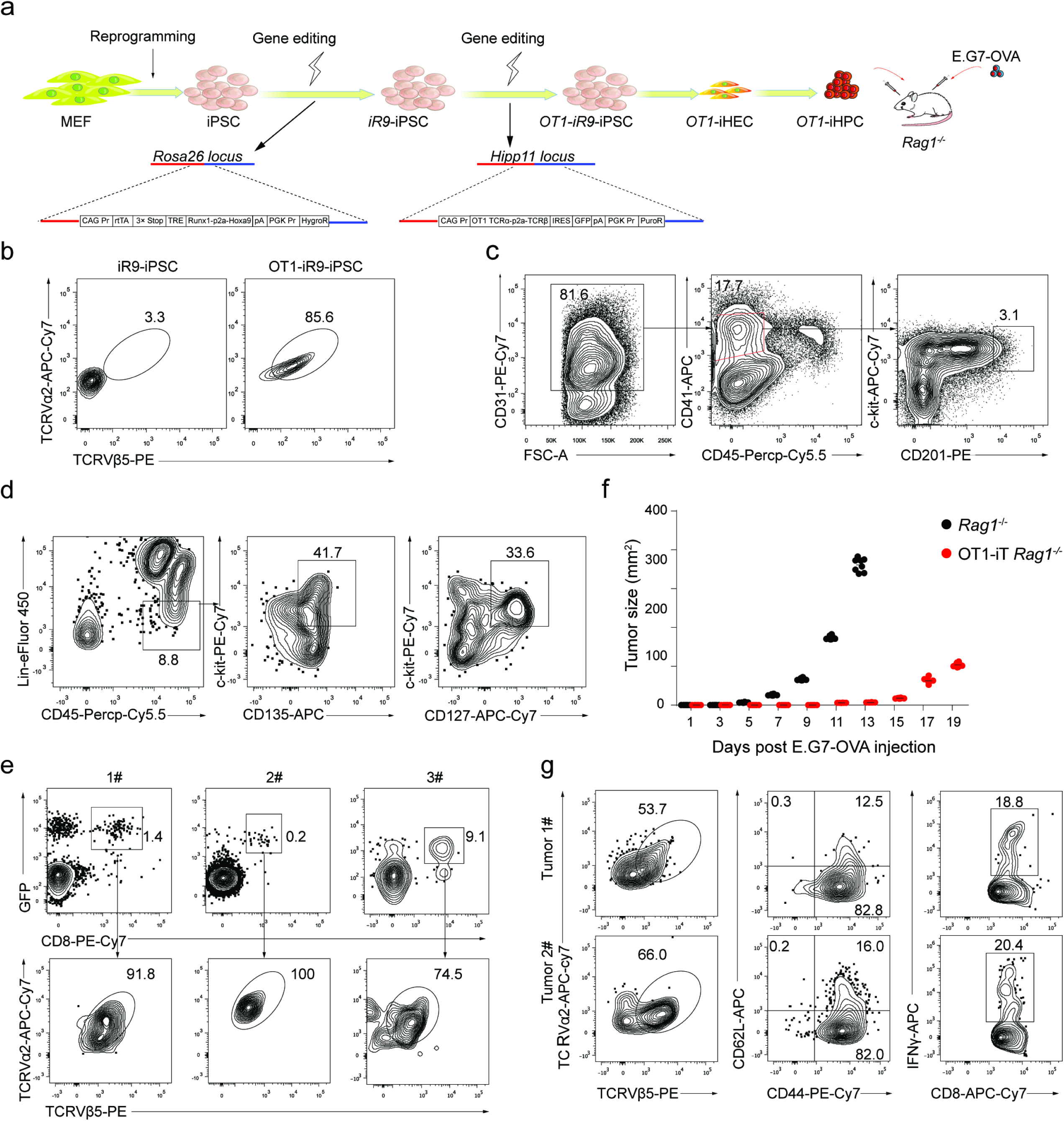
OT1-iT cell therapy suppresses the solid tumor growth in mice transplanted with E.G7-OVA cells. **a** Schematic diagram of OT1 engineered iT cells for anti-tumor therapy. Mouse MEF cells were isolated from CD45.2^+^ C57BL/6 mouse and reprogrammed into iPSC with Oct4, Klf4, and Sox2 retro-viruses. Then a *rtTA-TRE-Runx1-Hoxa9-HygroR* DNA cassette was inserted into the *Rosa26 locus*. Next, a *CAG-OT1-IRES-GFP-PuroR* expression element was inserted into the *Hipp11 locus* of *iR9*-iPSC. OT1-*iR9*-iPSC results in the production of CD8^+^ T cells carrying TCRVα2 and TCRVβ5 (MHC class I-restricted, ovalbumin-specific TCR). OT1-*iR9*-iPSC-derived iHEC were induced into iHPC (OT1-iHPC) as described in material and method sections. The iHPC were injected into irradiated (4.5 Gy) *Rag1^−/−^* recipient mice (3 million/mouse, 8-10-week-old C57BL/6 background). E.G7-OVA tumor cell line (C57BL/6 background) were transplanted into the groin of the *Rag1^−/−^* (n = 8) or OT1-iT-*Rag1^−/−^* (n = 8) by subcutaneous injection (0.2 million/mouse) six weeks after OT1-iHPC transplantation. **b** TCRVα2 and TCRVβ5 expression in OT1-*iR9*-iPSC measured by intracellular staining. The *iR9*-iPSC was used as negative control. **c** Sorting gates of the *OT1*-*iR9*-iPSC-derived iHEC population at day 11. The cells were enriched by streptavidin-beads recognizing biotin-CD31 before sorting. Representative plots from three independent experiments are shown. **d** Immuno-phenotypes of pre-thymic progenitors in induced hematopoietic progenitor cells from *OT1*-*iR9*-iPSC-derived iHEC after ten-day maturation. Representative plots from three independent experiments are shown. Lin was defined as CD2^−^CD3^−^CD4^−^CD8^−^CD11b^−^Gr1^−^Ter119^−^CD19^−^NK1.1^−^TCRγδ^−^. pre-thymic progenitors were defined as Lin^−^c-kit^+^CD127^+^/CD135^+^. **e** TCRVα2 and TCRVβ5 expression of iT cells in PB of *Rag1^−/−^* mice 6 weeks after transplantation of *OT1*-*iR9*-iPSC-derived iHPC. Three representative mice from three independent experiments were analyzed. **f** Tumor growth in *Rag1^−/−^* and OT1-iT-*Rag1^−/−^* mice. E.G7-OVA cells were transplanted into the groin of the *Rag1^−/−^* (n = 8) or OT1-iT-*Rag1^−/−^* mice (n = 8) by subcutaneous injection (0.2 million/mouse). The length and width of the tumors were measured every other day by a caliper, and each tumor size was calculated as length × width (mm^2^). Mice with tumor size larger than 20 mm at the longest axis were euthanized for ethical consideration. *** P < 0.001 (independent t-test, two-tailed). **g** Characterization of the OT1-iT cells in the tumors. The tumors were isolated at day 19 after injection and disaggregated by collagenase IV to single cell suspensions. The effector iT cells were defined as CD44^+^CD62L^−^. The memory iT cells were defined as CD44^+^CD62L^+^. IFNγ secreted by CD8^+^ OT1-iT cells in the tumors were intra-cellular stained. Representative plots from two tumors are shown.

## DISCUSSION

In this study, the iHEC from *iR9*-PSC gave rise to blood progenitor cells preferentially differentiating into iT cells *in vivo*. It is possible that the combinatory expression of *Runx1* and *Hoxa9*, pivotal transcription factors for definitive hematopoiesis ^22–24, 63^ and T cell development ^18^, synergistically and preferentially orchestrates the T and NK lineage potentials but intrinsically compromises the other blood lineage potentials during the early EHT and subsequent hematopoietic maturation phases in our induction protocol. Regarding the developmental evidence that an earlier wave of hematopoiesis preceding HSC emergence also produces blood progenitors possessing the T cell lineage potential ^14–16^, it is also possible that the *iR9*-PSC-derived iHPC resemble the developmental HPC prior to the occurrence of definitive HSC since overexpression of *Runx1* and *Hoxa9* at definitive HSC phase promoted myeloid-instead of lymphoid-biased hematopoiesis *in vivo* (Supplementary information, Fig. S8). The hematopoietic maturation step in the presence of OP9-DL1 feeder line unlikely causes T-lineage-biased iHPC, as an inducible expression of another transcription factor cocktail in PSC exactly using the same protocol gave rise to iHPC preferentially contributing to B lymphopoiesis in B-NDG recipients (unpublished data). Nonetheless, our data support the concept that synergies of distinct transcription factors intrinsically determine variable hematopoietic lineage potentials at as early as hemogenic endothelial cell stage.

Intravenous infusion of the iHPC from *iR9*-PSC successfully reconstituted iT lymphopoiesis *in vivo*. The induced LSK cells from the primary iHPC recipients further gave rise to T lymphocytes in secondary recipients. The occurrences of iDN1, iDN2, iDN3, iDN4 cells at different time-points in the thymi of iT-B-NDG mice strongly indicated that the induced pre-thymic progenitors (Lin^−^c-kit^+^CD127^+^/CD135^+^) have the capacities of homing to central lymphoid organs and developed normally following a cellular trajectory resembling natural T cell development. Despite the inefficient generation of CD4SP iT cells *in vitro* due to the MHC-I restricted OP9-DL1 feeder cells, robust phenotypic CD4SP iT cells generated *in vivo* and successful allogeneic rejection mediated by the CD4SP and CD8SP iT cells support that the regenerated regulatory iT cells possess normal immune functions. In combination with the new method of generating induced B (iB) cells (unpublished data), it would be promising to further test the coordinated immune responses of iT cells and iB cells in infection models. Besides the pivotal roles of exogenous *Runx1* and *Hoxa9* during EHT and subsequent iHPC maturation phases, we could not exclude the possibilities that the weak leaky expression of these two factors further facilitated the iT cell development *in vivo* after infusion into immune-deficient mice, as *Runx1* and *Hoxa9* are also involved in T cell development in bone marrow ^64^ and thymus ^18^. In contrast to our approach, an induced T cell progenitor population (DN2/DN3 cell phase) from mouse ESC lacked thymus-homing capacity *in vivo* and required congenic fetal thymus organ for further development into mature T cells ^6^, which implicated that an intrinsic gene network program essential for physiological T cell development were not fully activated during hematopoietic induction from PSC, which can be rescued by exogenous expression of *Runx1* and *Hoxa9*. Nonetheless, our approach fully reconstitutes functional T lymphopoiesis *in vivo* using PSC source, which avoids the malfunction risks of *in vitro* generated T cells due to the insufficiency of negative and positive selections.

The single iHEC exhibited a transcriptome signature resembling E11 AGM EC and pre-HSC. Activating the signature genes lacking in the iHEC but abundant in natural E11 AGM EC or pre-HSC might further promote the production of a homogenous iHEC population, thus consequently resulting in more efficient T cell generation or multi-lineage hematopoietic reconstitution. The feature of T cell-lineage-bias commitment from *iR9*-PSC brings advantages for gene editing using *iR9*-PSC rather than using canonical adult HSPC, since manipulating HSPC *in vitro* always faces stemness loss and might even introduce unknown impacts on the functions of other blood lineage derivatives from the edited HSPC.

In conclusion, this study establishes a novel approach of preferentially reconstituting functional and therapeutic T lymphopoiesis *in vivo* using PSC source by defined transcription factors. At single cell resolution, we unveil that the T-lineage specification is determined at as early as hemogenic endothelial cell stage and identify the *bona fide* pre-thymic progenitors with thymus-homing features. Given the enormous demand of regenerative T lymphopoiesis in treating T cell deficient and cancer-bearing patients, this study provides insight into therapeutic T lymphopoiesis using PSC source.

## ACKNOWLEDGEMENTS

This work is supported by the Strategic Priority Research Program of Chinese Academy of Sciences (XDA16010601), the CAS Key Research Program of Frontier Sciences (QYZDB-SSW-SMC057), the Major Research and Development Project of Guangzhou Regenerative Medicine and Health Guangdong Laboratory (2018GZR110104006), the Chinese Ministry of Science and Technology (2015CB964401, 2016YFA0100601, 2017YFA0103401, 2015CB964902, and 2015CB964901), the Health and Medical Care Collaborative Innovation Program of Guangzhou Scientific and Technology (201803040017), the Major Scientific and Technological Project of Guangdong Province (2014B020225005), the Science and Technology Planning Project of Guangdong Province (2017B030314056), the Program for Guangdong Introducing Innovative and Entrepreneurial Teams (2017ZT07S347), the National Natural Science Foundation of China (31471117, 81470281, 31600948, 31425012).

## AUTHOR CONTRIBUTIONS

R.G. and F.H. conducted all the major experiments, data analysis and wrote the manuscript. Q.W., C.Lv, H.W., L.L., Y.Z., Z.B., M.Z., Y.L., X.L., C.X., T.W., P.Z., K.W., Y.D., Y.L., YX.G. and Y.G. participated in multiple experiments; Q.W. and Z.L. performed RNA-Seq and data analysis. C.Lv, H.W., Y.L., P.Z., Y.L., X.Z. and J.C. constructed vectors, prepared iPSC, designed and participated gene editing. Y.L. and YQ.L. discussed the single cell data; B.L. and J.W. discussed the data and wrote the manuscript; and J.W. designed the project and provided the final approval of the manuscript.

## MATERIALS AND METHODS

### Mice

B-NDG (NOD-*Prkdc^Scid^IL2rg^tm^*^1^/Bcgen, CD45.1^+^) mice were purchased from Biocytogen Jiangsu Co., Ltd (Jiangsu, China). BALB/c and C57BL/6 (CD45.2^+^) mice were purchased from Bejing Vital River Laboratory Animal Technology. *Rag1^−/−^* mice (C57BL/6 background) were a gift from Dr. Z. Liu from the Institute of Biophysics (CAS, China). Mice were housed in the SPF-grade animal facility of the Guangzhou Institutes of Biomedicine and Health, Chinese Academy of Sciences (GIBH, CAS, China). All animal experiments were approved by the Institutional Animal Care and Use Committee of Guangzhou Institutes of Biomedicine and Health (IACUC-GIBH).

### Cell culture

Mouse embryonic fibroblasts (MEFs) were derived from 13.5 d.p.c C57BL/6 mouse embryos. MEFs were maintained in DMEM/high glucose (Hyclone), 10% FBS (Natocor) supplemented with 1% nonessential amino acids (NEAA, Gibco). C57BL/6 mouse embryonic stem cells (Biocytogen) were maintained on feeder layers in ES medium containing DMEM/high glucose, 15% FBS (Gibco), 1% NEAA, 1% GlutaMAX (Gibco), 1% Sodium Pyruvate (Gibco), 0.1 mM β-mercaptoethanol (Gibco), 1 μM PD0325901 (Selleck), 3 μM Chir99021 (Selleck) and 1000 U/ml LIF. The OP9-DL1 cells (GFP^+^) were maintained in α-MEM (Gibco) supplemented with 20% FBS (CellMax). The AFT024 cell lines (ATCC) were maintained in DMEM/high glucose, 10% FBS (Natocor) supplemented with 0.1 mM β-mercaptoethanol and 1% Sodium Pyruvate. HEK293T (ATCC) and Plat-E (Cell Biolabs, Inc) cells were maintained in DMEM/high glucose supplemented with 10% FBS (Natocor). E.G7-OVA cell line (ATCC) was cultured in RPMI 1640 (Gibco) supplemented with 10% FBS (Natocor), 1% GlutaMAX, 1% sodium pyruvate, and 0.1 mM β-mercaptoethanol.

### Hematopoietic differentiation

PSC were trypsinized by 0.05% Trypsin-EDTA (Gibco) and resuspended in the basic differentiation medium (BDM: IMDM, 15% FBS (Gibco), 200μg/ml iron-saturated transferring (Sigma), 0.45 mM monothiolglycerol (Sigma), 1% GlutaMAX, and 50 μg/ml ascorbic acid (Sigma)). For removing the feeder layers, the PSC were plated into the 0.1% gelatin-coated (Merck Millipore) well, and the floating cells were collected after 40 min. For EB generation ^65^, the PSC were resuspended at 100,000 cells/ml in the BDM supplemented with 5 ng/ml BMP4 (Peprotech) and plated at 20 ul/drop for inverted culture in 15 cm dishes. At day 2.5, EBs were replanted into gelatinized plates in BDM supplemented with 5 ng/ml BMP4 and 5 ng/ml VEGF (Peprotech). At day 6, the medium was changed to BDM supplemented with 2% conditioned medium derived from the supernatants of AFT024-mIL3, AFT024-mIL6, AFT024-hFlt3L and AFT024-mSCF cell culture. Doxycycline (1 μg/ml, Sigma) was added at day 6. The medium was replaced every other day. The plates were seeded with OP9-DL1 cells (20000 cells/well, 12-well plate) 12 hours prior to the hematopoietic maturation step in EM medium (α-MEM, 15% DFBS (Hyclone), 200 μg/ml iron-saturated transferring, 0.45 mM monothiolglycerol, 1% GlutaMAX, 50 μg/ml ascorbic acid, 2% conditioned medium derived from supernatants of AFT024-mIL3, AFT024-hFlt3L and AFT024-mSCF cell culture and 1 μg/ml doxycycline. 100-500 sorted iHEC were seeded into each well for hematopoietic maturation. The EM was half-replaced every two days.

### Transplantation of iHPC

8-10-week-old B-NDG mice were sublethally irradiated (2.25 Gy) by an X-ray irradiator (RS2000, Rad Source Inc.). 0.5-1 million PSC-derived iHPC were injected into each irradiated B-NDG mouse via retro-orbital veins. The mice were fed with water containing co-trimoxazole (Tianjin Lisheng Pharmaceutical co., LTD) for two weeks to prevent infection.

### T lymphocyte induction *in vitro*

For T lymphocyte induction *in vitro*, OP9-DL1 coculture method^1^ was used with minor modifications. Briefly, the single-cell suspensions of iHPC (Day 21) were maintained on OP9-DL1 feeder cells in T cell induction medium (TIM, α-MEM, 20% DFBS, and 1% GlutaMAX) supplemented with 2% conditioned medium derived from supernatants of AFT024-hFlt3L and AFT024-hIL7 cell culture for sustained 12 days. The iHEC-derived cells were trypsinized into single-cell suspensions and replanted into fresh OP9-DL1 monolayers every 6 days. And the TIM was replaced every 3 days.

### Gene editing

Mouse MEF cells were reprogrammed into iPSC as described ^66^. The *CAG Pr-rtTA-3×Stop-TRE-Runx1-p2a-Hoxa9-pA-PGK Pr-HygroR* cassette was inserted into the *Rosa26 locus* of mouse ESC/iPSC. The positive clones (*iR9*-ESC/iPSC) were selected by Hygromycin B (150 μg/ml, Invivogen) were further cultured in ES medium supplemented with Dox (1 μg/ml). The induced expression of *Runx1* and *Hoxa9* was confirmed by qPCR. For the generation of *OT1*-*iR9*-iPSC, a *CAG Pr-OT1 TCRα-p2a-TCRβ-IRES-GFP-PGK Pr-PuroR* cassette was inserted into the *Hipp11 locus* of *iR9*-iPSC. The *OT1* sequence was cloned from murine TCR OT1-2A.pMIG II (Addgene). The *OT1*-*iR9*-iPSC positive clones were further selected by Puromycin (1 μg/ml, Invivogen) and the expression of OT1-TCR were measured by intra-cellular staining.

### Flow cytometry and cell sorting

Single-cell suspensions were prepared by 0.05% Trypsin-EDTA and filtered by 70 μm filter. Single cells were blocked by Fc (CD16/32) (93, eBioscience) antibody, and then stained with related antibodies. The following antibodies were used: c-kit (2B8, eBioscience), CD31 (390, eBioscience), CD41 (eBioMWReg30, eBioscience), CD45 (30-F11, eBioscience), CD45.1 (A20, eBioscience), CD45.2 (104, eBioscience), CD2 (RM2-5, eBioscience), CD3 (145-2C11, eBioscience), CD4 (GK1.5, eBioscience), CD8a (53-6.7, eBioscience), CD19(eBio1D3, eBioscience), B220 (RA3-6B2, eBioscience), CD11b (M1/70, eBioscience), NK1.1 (PK136, eBioscience), Ter119 (TER-119, eBioscience), Gr1 (RB6-8C5, eBioscience), CD201 (eBio1560, eBioscience), CD135 (A2F10, eBioscience), CD127 (A7R34 eBioscience) FcεRIα (MAR-1, biolegend), CD69 (H1.2F3, biolegend), CD62L (MEL-14, biolegend) IFNγ (XMG1.2, biolegend), IL17 (TC11-18H10.1, biolegend), CD44 (IM7, eBioscience), CD25 (PC61.5, eBioscience), TCRβ (H57-597, eBioscience), TCRγδ (GL3, eBioscience), TCRvα2 (B20.1, biolegend), TCRvβ5.1/5.2 (MR9-4, biolegend) Streptavidin PE-Cy7 (eBioscience), Streptavidin eFlour 450 (eBioscience), Streptavidin PE-Cy5 (biolegend). The cells were resuspended in the DAPI solution, or PI solution (eBioscience) and were analyzed with Fortessa cytometer (BD Biosciences). The cells were sorted using Arial II cytometer (BD Biosciences). The flow data were analyzed with FlowJo (Three Star, Ashland OR).

### Allogeneic skin transplantation

Individual *Rag1^−/−^* mice (8-10 weeks old) were adoptively transferred with splenic cells equivalent to 5 million CD4^+^ and CD8^+^ iT cells from iT-B-NDG mice. Four days after iT cell transfer, the allogeneic skin (BALB/c background) was transplanted as described ^67^. Grafts were considered rejection if there was a loss of distinct border, visible signs of ulceration and necrosis to 80% of the graft area. The rejected skin tissues were removed for analysis 9 days after skin transplantation. For analysis activated iT cells in rejected skin grafts, the single cell suspensions were prepared as described ^68^. The activated alloreactive iT lymphocytes were defined as CD45.2^+^Ter119^−^CD11b^−^CD69^+^CD44^+^CD4^+^/CD8^+^. For analysis of cytokines released by the alloreactive iT cells, we used anti-IL17 and anti-IFNγ antibodies following an intracellular staining protocol (eBioscience).

### OT1-iT anti-tumor assay

For the reconstitution of the OT1-iT cells in *Rag1^−/−^* mice, three million OT1-iHPC were transplanted into each irradiated *Rag1^−/−^* mouse (4.5 Gy). OT1-iT cells (GFP^+^ CD8^+^ TCRVβ5^+^ TCRVα2^+^) in PB were analyzed six weeks post-transplantation. The E.G7-OVA cells were transplanted into the groin of the OT1-iT reconstituted mice by subcutaneous injection (0.2 million/mouse). The tumor size was measured every 2 days and was calculated as length × width (mm^2^). Mice with tumor size larger than 20 mm at the longest axis were euthanized for ethical consideration. To analyze the tumor-infiltrating OT1-iT cells, tumors were isolated at day 15 and digested for 30 min at 37 < by collagen IV solution (1mg/ml, Gibco) after being cut up. Then, the single-cell suspensions were harvested for staining. The activated iT cells were defined as CD45.2^+^GFP^+^CD8^+^CD44^+^CD62L^−^.

### RNA-seq and data analysis

The cDNA of single iHEC sorted on day 11, and iHPC at Day 14, 17, and 21 or 1,000-CD4SP/CD8SP iT-cell aliquots of from spleens of iT-B-NDG mice were generated and amplified using Discover-sc WTA Kit V2 (Vazyme). The quality of amplified cDNA was assessed by qPCR analysis of housekeeping genes (B2m and Gapdh). Samples that passed quality control were used for sequencing library preparation by TruePrep DNA Library Prep Kit V2 (Vazyme). All libraries were sequenced by illumina sequencer NextSeq 500. The raw data (fastq files) were generated using bcl2fastq software (version 2.16.0.10) and were uploaded to the Gene Expression Omnibus public database (GSE121371, GSE121373, GSE128738). The raw reads were aligned to mouse genome mm10 by HISAT2 (version 2.1.0) ^69^ and the expression levels in TPM were estimated by StringTie (version 1.3.4) ^70, 71^. The wildtype CD4SP T cells, CD8SP T cells, myeloid cells, and B cells sequencing data (GSE105057) were downloaded from Gene Expression Omnibus ^13^. Heat maps were plotted using pheatmap (version 1.0.8). The natural embryonic single-cell data (endothelial cells (CD31^+^VE-cadherin^+^CD41^−^CD43^−^CD45^−^Ter119^−^) T1 pre-HSC (CD31^+^CD45^−^CD41^low^c-kit^+^CD201^high^), T2 pre-HSC (CD31^+^CD45^+^c-Kit^+^CD201^+^), E12 HSC (Lin^−^Sca-1^+^CD11b^low^CD201^+^), E14 HSC (CD45^+^CD150^+^CD48^−^CD201^+^), and adult HSC (CD45^+^CD150^+^CD48^−^CD201^+^)) were downloaded from Gene Expression Omnibus (GSE67120) ^35^.The batch effects of single-cell data between iHEC and natural embryonic cells were removed using ComBat (sva R package, version 3.26.0). The prcomp function of stats (R package, version 3.4.4) was used for PCA. The DESeq2 was used for differential expression analysis. The PCA plot and violin plot were plotted using ggplot2 (R package, version 2.2.1). tSNE was performed by Rtsne (R package version 0.15). The TPM values of transcription factors were log2-converted.

For TCRαβ sequencing, 15,000 sorted CD4SP, and CD8SP naïve iT cells were sorted from thymus or spleen of iT-B-NDG mice. The sorted iT cells of thymus were gated on CD45.2^+^Ter119^−^CD11b^−^Gr1^−^CD19^−^B220^−^NK1.1^−^TCRγδ^−^CD4^+^CD8^−^ and CD45.2^+^Ter119^−^CD11b^−^Gr1^−^CD19^−^B220^−^NK1.1^−^TCRγδ^−^CD4^−^CD8^+^. The splenic naïve iT cells were gated on CD45.2^+^CD4^+^CD8^−^CD62L^+^CD44^−^ and CD45.2^+^CD4^−^CD8^+^CD62L^+^CD44^−^. The cDNA was generated and amplified by SMARTer Mouse TCRαβ Profiling Kit (Clontech). Libraries were sequenced by illumina sequencer MiSeq (2×250 cycles). The raw data (fastq files) were generated using illumina bcl2fastq software and were uploaded to Gene Expression Omnibus public database (GSE121374). T cell receptors αβ chains repertoires were aligned and assembled using software MiXCR (version 2.1.12) ^45^. The TCRαβ clonotypes were exported respectively by parameter ‘--chains’ in exportClones command of MiXCR. The exported clonotypes were visualized in the form of chord diagram using VDJtools software (version 1.1.10) ^72^.

### Statistics

All quantitative analyses were based on a minimum of at least three sample replicates. Data are presented as means ± s.d. by GraphPad Prism. Independent-sample student T test and One-way ANOVA were performed (SPSS). NS, not significant; *p < 0.05; **p < 0.01; ***p < 0.001.

