## Supplementary information, Fig. S1 for "Guiding T lymphopoiesis from pluripotent stem cells by defined transcription factors"

Figure S1

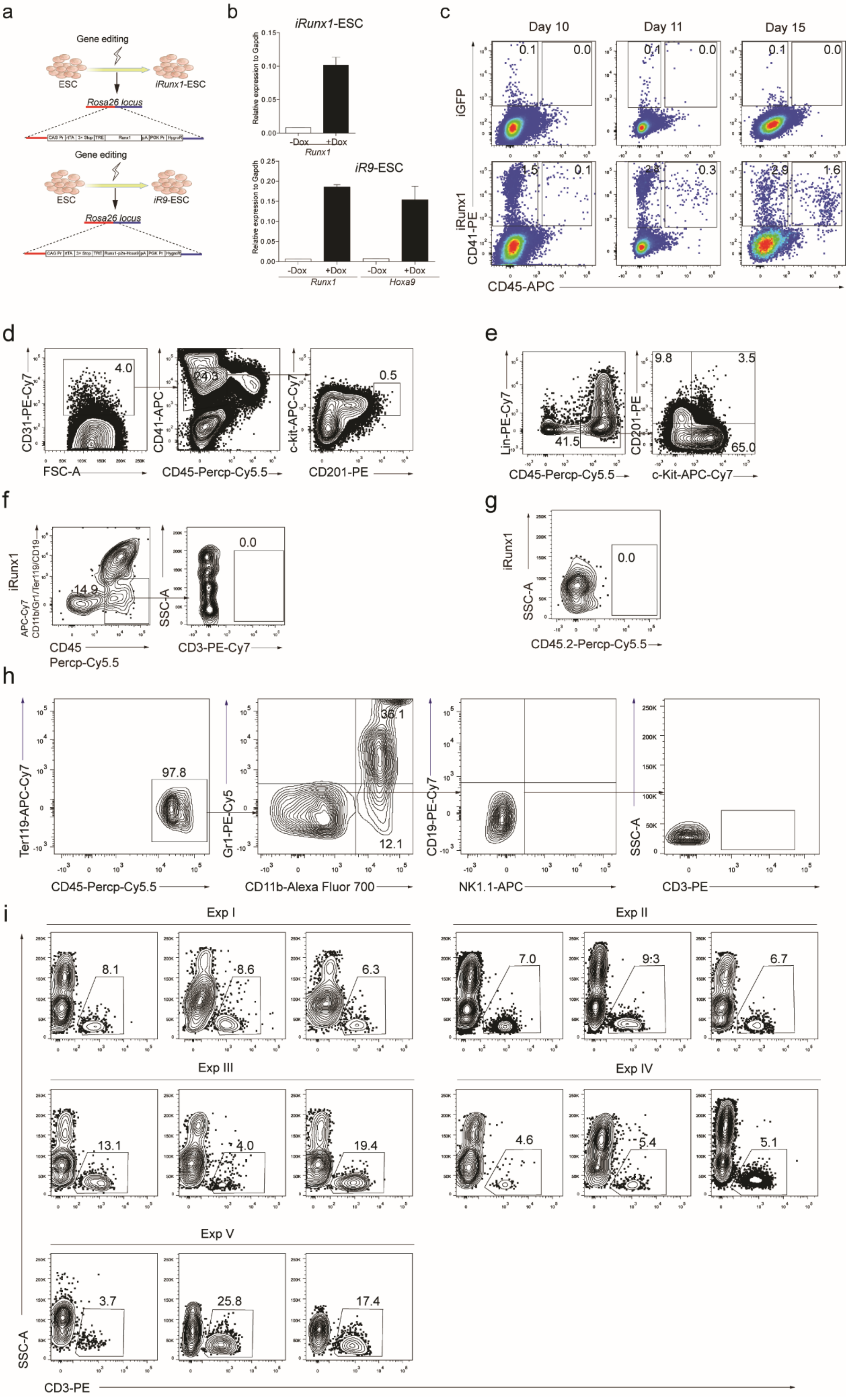
a
Gene editing
ESC
iRunx1-ESC
Rosa26 locus
Gene editing
ESC
iR9-ESC
Rosa26 locus
b
iRunx1-ESC
Relative expression to Gapdh
-Dox
+Dox
Runx1
iR9-ESC
Relative expression to Gapdh
-Dox
+Dox
-Dox
+Dox
Runx1
Hoxa9
c
Day 10
Day 11
Day 15
iGFP
iRunx1
CD41-PE
CD45-APC
d
CD31-PE-Cy7
FSC-A
4.0
CD41-APC
CD45-Percp-Cy5.5
24.5
c-kit-APC-Cy7
CD201-PE
0.5
e
Lin-PE-Cy7
CD45-Percp-Cy5.5
41.5
CD201-PE
c-Kit-APC-Cy7
9.8
3.5
65.0
f
iRunx1
APC-Cy7
CD11b/Gr1/Ter119/CD19
CD45
Percp-Cy5.5
14.9
SSC-A
CD3-PE-Cy7
0.0
g
iRunx1
SSC-A
CD45.2-Percp-Cy5.5
0.0
h
Ter119-APC-Cy7
CD45-Percp-Cy5.5
97.8
Gr1-PE-Cy5
CD11b-Alexa Fluor 700
36.1
12.1
CD19-PE-Cy7
NK1.1-APC
SSC-A
CD3-PE
i
Exp I
8.1
8.6
6.3
Exp II
7.0
9.3
6.7
Exp III
13.1
4.0
19.4
Exp IV
4.6
5.4
5.1
Exp V
3.7
25.8
17.4
SSC-A
CD3-PE

**Supplementary Figure 1.** Transcription factor Runx1 as the starting candidate for iT cell regeneration. **a** Schematic diagram of iRunx1-ESC and iRunx1-p2a-Hoxa9 (*iR9*)-ESC construction. A rtTA-TRE-Runx1-HygroR DNA cassette or rtTA-TRE-Runx1-Hoxa9-HygroR DNA cassette was inserted into the *Rosa26 locus* of CD45.2$^{+}$ C57BL/6 mouse ESC by homologous recombination. The *iRunx1*- or *iR9*-ESC clones were selected by hygromycin B (150 μg/ml). **b** Q-PCR analysis of the expression of Runx1 in iRunx1-ESC, and the expression of Runx1 and Hoxa9 in *iR9*-ESC after doxycycline induction. **c** Runx1 expression promotes endothelial to hematopoietic transition. CD41$^{+}$CD45$^{-}$ and CD41$^{+}$CD45$^{+}$ populations of iGFP-ESC-derived and iRunx1-ESC-derived cells were analyzed at day 10, day 11 and day 15 by flow cytometry. **d** Sorting gates of iRunx1 ESC-derived iHEC population at day 11. Representative plots from three independent experiments are shown. **e** Immuno-phenotypes of hematopoietic progenitors from iRunx-ESC-derived iHEC after ten-day maturation. Representative plots from four independent experiments are shown. Lin was defined as CD2$^{-}$CD3$^{-}$CD4$^{-}$CD8$^{-}$Gr1$^{-}$Ter119$^{-}$CD19$^{-}$NK1.1$^{-}$TCRγδ$^{-}$. Hematopoietic progenitors were defined as CD45$^{+}$Lin$^{-}$c-kit$^{+}$CD201$^{+}$. **f** The failure of iT cell induction from iRunx1-ESC-derived iHEC. Flow cytometry analysis of induced T cells from in vitro induction of iRunx1-ESC-derived iHEC. One thousand cells of iRunx1-ESC-derived iHEC were seeded into per well (24-well plates) pre-seeded with OP9-DL1 feeder cells (10000 cells/well) 12 hours prior maturation in EM medium with doxycycline (1 μg/ml). Doxycycline was sustained for 10 days during the maturation step. After maturation, the bulk blood cells were assessed for T lineage generation potential. For in vitro T cell induction, the medium was changed to T cell induction medium (TIM, α-MEM, 20% DFBS, and 1% GlutaMAX) supplemented with 2% conditioned medium derived from supernatants of AFT024-hFlt3L and AFT024-hIL7 cell culture for sustaining 12 days. **g** iRunx1-ESC-derived iHEC failed to contribute any hematopoietic in B-NDG mice four weeks after transplantation. One million iHEC-derived hematopoietic cells were transplanted into individual B-NDG mice (CD45.1$^{+}$) irradiated by X-ray (2.25 Gy). One representative mouse was analyzed. **h** Lineage characterization of bulk cells collected at day 10 after coculture of *iR9*-ESC-derived iHEC and OP9-DL1. One representative plot from three independent experiments are shown. Myeloid cells were defined as CD45$^{+}$CD11b$^{+}$. **i** Flow cytometry analysis of pluripotent stem cell-derived T cells in PB of iHPC recipients. Plots from three mice of each independent experiment (Exp I, II, III, IV, V) are shown.
