## Supplementary information, Fig. S2 for "Guiding T lymphopoiesis from pluripotent stem cells by defined transcription factors"

Figure S2

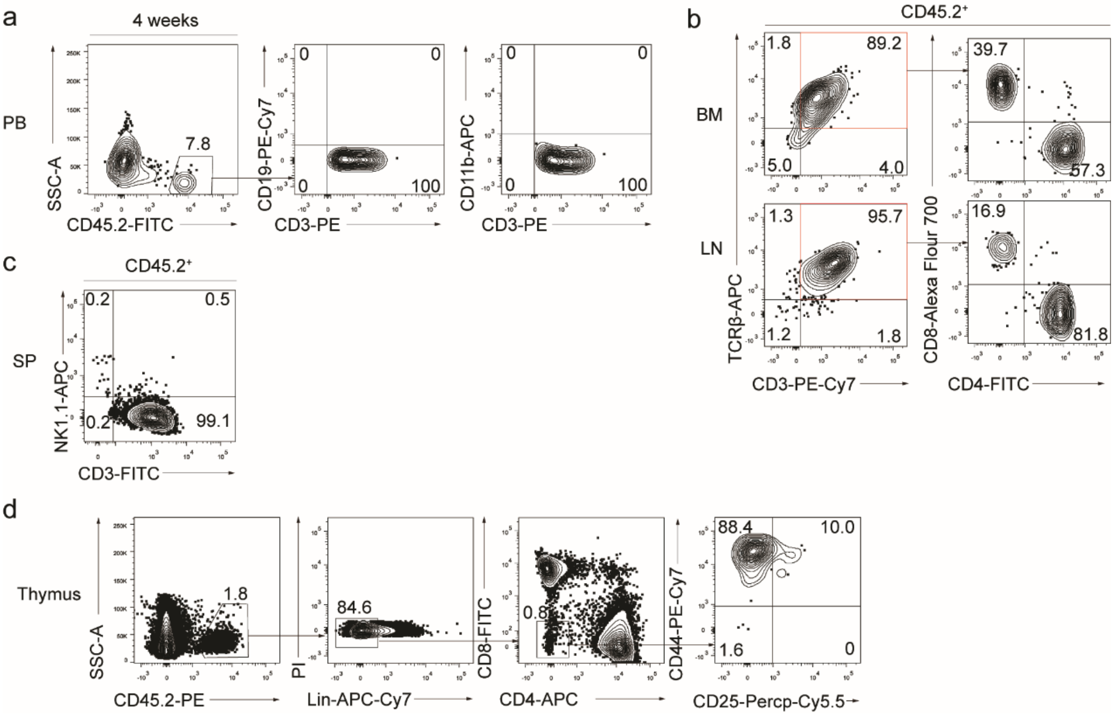

a
4 weeks
PB
SSC-A
CD45.2-FITC
7.8
CD19-PE-Cy7
CD3-PE
0
0
0
100
CD11b-APC
CD3-PE
0
0
0
100
b
CD45.2+
BM
LN
1.8
89.2
5.0
4.0
1.3
95.7
1.2
1.8
TCRβ-APC
CD3-PE-Cy7
39.7
57.3
16.9
81.8
CD8-Alexa Flour 700
CD4-FITC
c
CD45.2+
SP
NK1.1-APC
CD3-FITC
0.2
0.5
0.2
99.1
d
Thymus
SSC-A
CD45.2-PE
1.8
PI
Lin-APC-Cy7
84.6
CD8-FITC
CD4-APC
0.8
CD44-PE-Cy7
CD25-Percp-Cy5.5
88.4
10.0
1.6
0

**Supplementary Figure 2.** Day-17 iHPC reconstitutes T lymphopoiesis *in vivo.* **a** Induced T cells (iT) in PB of B-NDG mice were analyzed by flow cytometry 4 weeks after transplantation. One million iHPC from Day 17 in the presence of OP9-DL1 feeder cells were transplanted into individual B-NDG mice ($CD45.1^+$) irradiated by X-ray (2.25 Gy). One representative mouse from three independent experiments were analyzed. **b** Flow cytometry analysis of the mature iT cells in bone marrow (BM) and lymph node (LN) of B-NDG mice transplanted with iHPC. Each B-NDG mouse was transplanted with one million iHPC collected at day 6 in the presence of OP9-DL1 feeder cells. Representative mouse was sacrificed and analyzed 5 weeks after transplantation. Data from one mouse are shown. **c** Flow cytometry analysis of induced NK (iNK) cells and induced T cells (iT) in spleen (SP) of B-NDG mice transplanted with iHPC. Each B-NDG mouse was transplanted with one million iHPC collected from day-6 co-culture with OP9-DL1. Representative mouse was sacrificed and analyzed 5 weeks after transplantation. iNK cells were defined as $CD45.2^+NK1.1^+CD3^-$. Data from one representative mouse were shown. **d** Flow cytometry analysis of induced double-negative T lymphocytes (iDN) cells in the thymus of B-NDG mice transplanted with iHPC. Each B-NDG mouse was transplanted with one million iHPC after six-day co-culture with OP9-DL1. Representative mouse was sacrificed and analyzed 5 weeks after transplantation. Data from one representative mouse from three independent experiments are shown. Lin was defined as $Ter119^-CD11b^-Gr1^-CD19^-B220^-NK1.1^-TCR\gamma\delta^-$.
