## Supplementary information, Fig. S3 for "Guiding T lymphopoiesis from pluripotent stem cells by defined transcription factors"

Figure S3

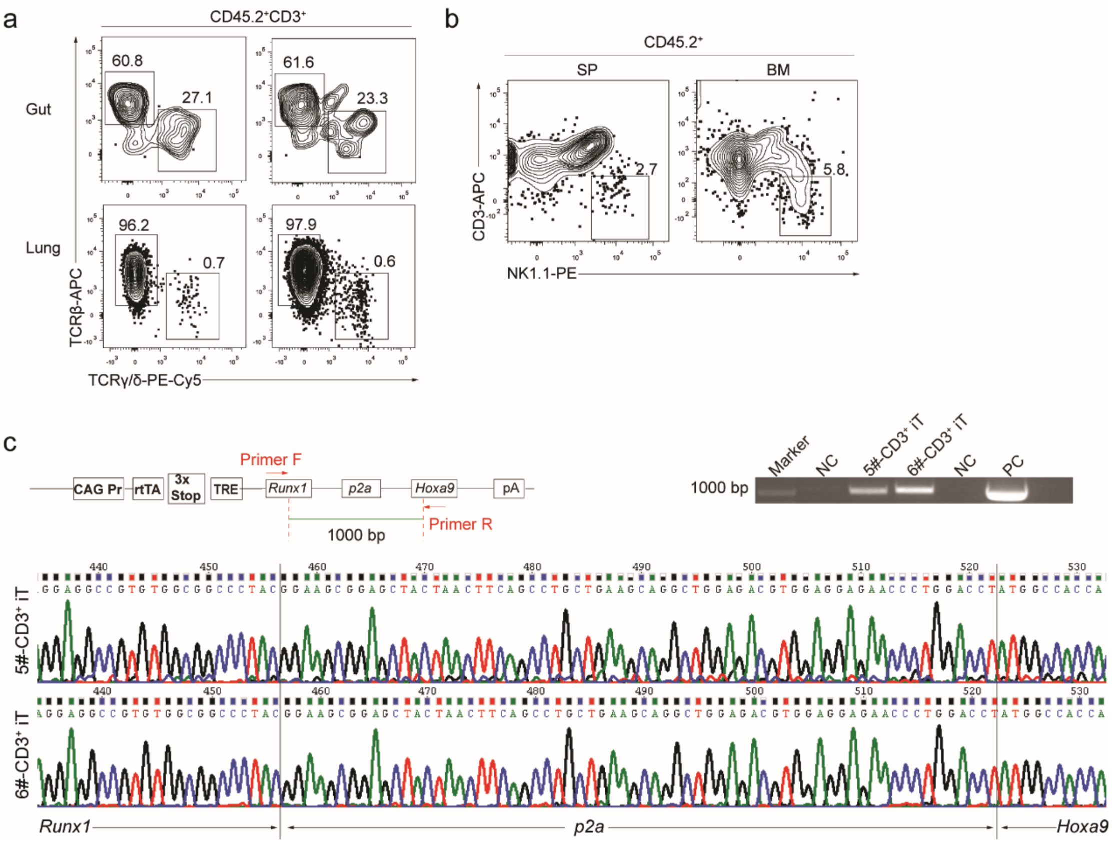

a
CD45.2⁺CD3⁺
Gut
Lung
60.8
27.1
61.6
23.3
96.2
0.7
97.9
0.6
TCRβ-APC
TCRγ/δ-PE-Cy5
b
CD45.2⁺
SP
BM
2.7
5.8
CD3-APC
NK1.1-PE
c
CAG Pr
rtTA
3x Stop
TRE
Primer F
Runx1
p2a
Hoxa9
pA
Primer R
1000 bp
Marker
NC
5#-CD3⁺ iT
6#-CD3⁺ iT
NC
PC
1000 bp
5#-CD3⁺ iT
6#-CD3⁺ iT
Runx1
p2a
Hoxa9

**Supplementary Figure 3.** Characterization of iHEC-derived iT and iNK cells in B-NDG recipients. **a** Flow cytometry analysis of γδ-iT and β-iT cells in the gut and lung tissues of B-NDG mice transplanted with ESC-derived hematopoietic cells. Each B-NDG mouse was transplanted with one million iHEC-derived hematopoietic cells and fed with Doxycycline (1 mg/ml) for sustaining 6 weeks. Representative mouse was sacrificed and analyzed at 6 weeks after transplantation. Data from two representative mice are shown. **b** Measurement of induced NK cells in spleen (SP) and bone marrow (BM) of B-NDG mice transplanted with iHPC. Pluripotent stem cell-derived NK cells were defined as $CD45.2^{+}NK1.1^{+}CD3^{-}$. Plots from one representative B-NDG mouse six weeks after transplantation of iHPC are shown. **c** Genotype sequencing of the Runx1-p2a-Hoxa9 element of the sorted iT cells. Genomic PCR were performed using primer pairs flanking the inserted Runx1-p2a-Hoxa9 sequence element, and genome template (200 ng) of 20,000 sorted iT cells from the spleen of iT-B-NDG mice. The sequencing results were visualized by chromas software. PCR products of iT cells from two iT-B-NDG mice were sequenced and shown.
