## Supplementary information, Fig. S4 for "Guiding T lymphopoiesis from pluripotent stem cells by defined transcription factors"

### Figure S4

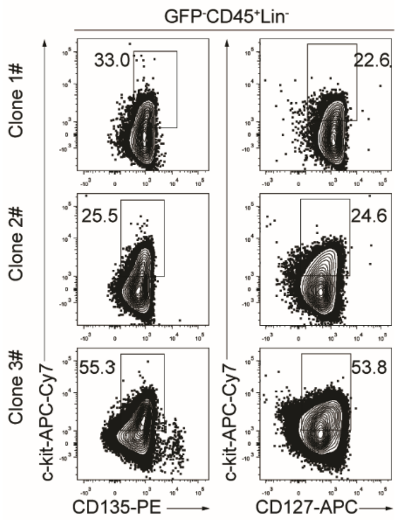
GFP-CD45+Lin-
Clone 1#
Clone 2#
Clone 3#
33.0
22.6
25.5
24.6
55.3
53.8
c-kit-APC-Cy7
c-kit-APC-Cy7
CD135-PE
CD127-APC

**Supplementary Figure 4.** Single iHEC from *iR9*-ESC gives rise to iHPC *in vitro.* Immuno-phenotypes of pre-thymic progenitors in induced hematopoietic progenitor cells from *iR9*-PSC-derived iHEC after ten-day maturation. Three representative clones from two independent experiments were analyzed. Lin was defined as $CD2^-CD3^-CD4^-CD8^-CD11b^-Gr1^-Ter119^-CD19^-NK1.1^-TCR\gamma\delta^-$. pre-thymic progenitors were defined as $Lin^-c\text{-}kit^+CD127^+/CD135^+$.
