## Supplementary information, Fig. S5 for "Guiding T lymphopoiesis from pluripotent stem cells by defined transcription factors"

Figure S5

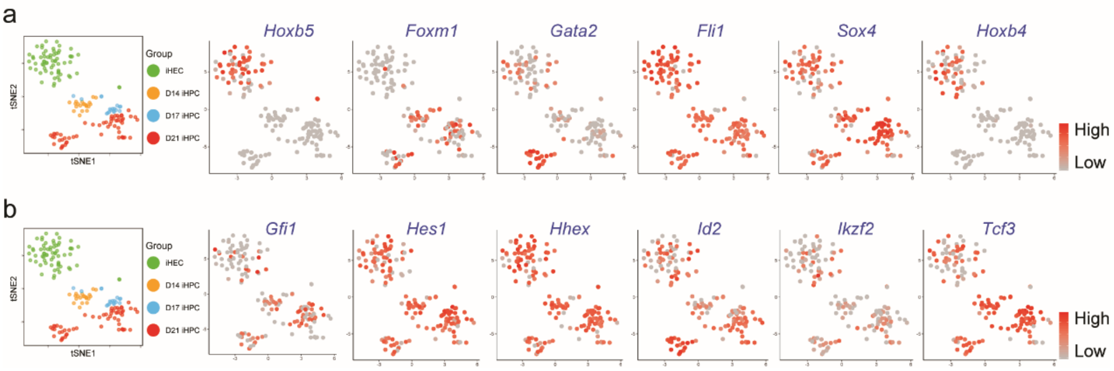
a
Group
iHEC
D14 iHPC
D17 iHPC
D21 iHPC
tSNE1
tSNE2
Hoxb5
Foxm1
Gata2
Fli1
Sox4
Hoxb4
High
Low
b
Gfi1
Hes1
Hhex
Id2
Ikzf2
Tcf3

**Supplementary Figure 5.** Time-course expression patterns of hematopoietic and lymphopoietic development-related transcription factors in single iHEC and iHPC derived from *iR9*-ESC. **a** tSNE analysis of the expression pattern of selected hematopoietic-related transcription factors (*Hoxb5*, *Foxm1*, *Gata2*, *Fli1*, *Sox4*, and *Hoxb4*) in iHEC and iHPC. **b** tSNE analysis of the expression pattern of selected T cell development-related transcription factors (*Gfi1*, *Hes1*, *Hhex*, *Id2*, *Ikzf2*, and *Tcf3*) in iHEC and iHPC at Day14, Day17, and Day21.
