## Supplementary information, Fig. S6 for "Guiding T lymphopoiesis from pluripotent stem cells by defined transcription factors"

### Figure S6

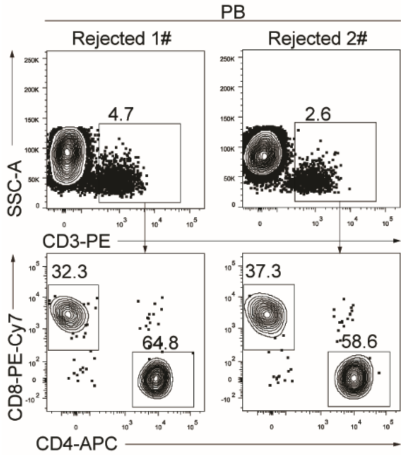
PB
Rejected 1#
Rejected 2#
SSC-A
CD3-PE
4.7
2.6
CD8-PE-Cy7
CD4-APC
32.3
64.8
37.3
58.6

**Supplementary Figure 6.** iT cells in PB of iT-*Rag1*$^{-/-}$ mice 30 days after 1$^{st}$ allogeneic rejection. Flow cytometry analysis of the iT cells in peripheral blood (PB) of iT transferred *Rag1*$^{-/-}$ recipients 30 days after 1$^{st}$ grafted allogeneic skin rejection. Plots of two representative mice are shown.
