## Supplementary information, Fig. S7 for "Guiding T lymphopoiesis from pluripotent stem cells by defined transcription factors"

### Figure S7

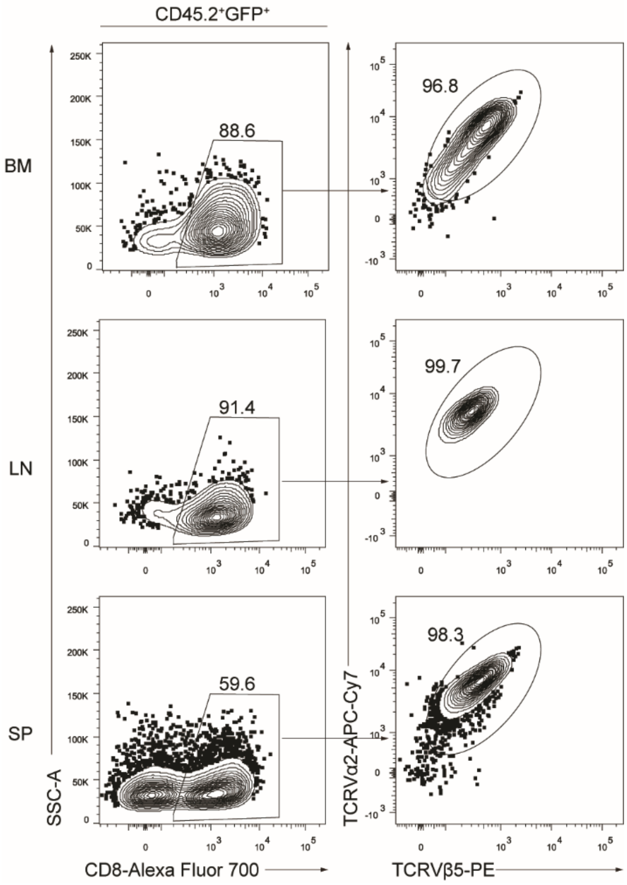
CD45.2+GFP+
BM
LN
SP
88.6
91.4
59.6
96.8
99.7
98.3
SSC-A
CD8-Alexa Fluor 700
TCRVα2-APC-Cy7
TCRVβ5-PE

**Supplementary Figure 7.** Distributions of OT1-iT cells in the BM, LN and SP of OT1-iT *Rag1*$^{-/-}$ mice 19 days after E.G7-OVA tumor cell injection. *Rag1*$^{-/-}$ mice were transplanted with iHPC (3 million/mouse) 4 weeks before E.G7-OVA tumor cell injection. OT1-iT *Rag1*$^{-/-}$ mice carrying tumors were sacrificed 19 days after E.G7-OVA tumor cell injection. TCRVα2 and TCRVβ5 on CD8$^{+}$ iT cells were analyzed in the bone marrow (BM), lymph node (LN) and spleen (SP) of the OT1-iT *Rag1*$^{-/-}$ mice. Plots of one representative mouse are shown.
