## Supplementary information, Fig. S8 for "Guiding T lymphopoiesis from pluripotent stem cells by defined transcription factors"

### Figure S8

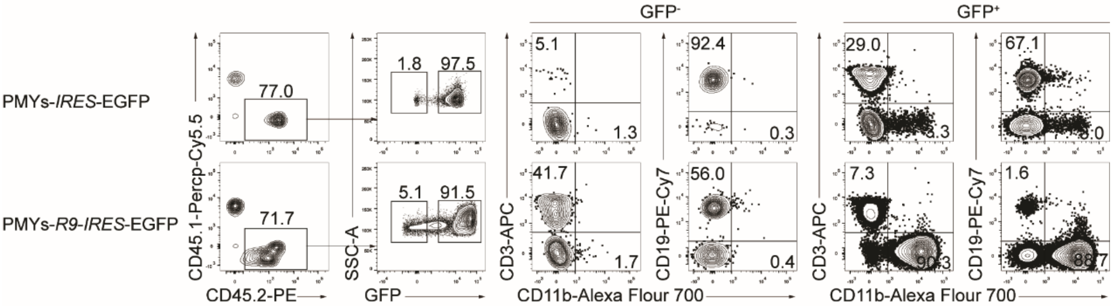

GFP⁻
GFP⁺
PMYs-IRES-EGFP
PMYs-R9-IRES-EGFP
77.0
71.7
1.8
97.5
5.1
91.5
5.1
1.3
41.7
1.7
92.4
0.3
56.0
0.4
29.0
7.3
1.6
67.1
CD45.1-Percp-Cy5.5
CD45.2-PE
SSC-A
GFP
CD3-APC
CD19-PE-Cy7
CD11b-Alexa Flour 700

**Supplementary Figure 8.** Overexpression of tandem *Runx1-p2a-Hoxa9* in natural HSC leads to myeloid- instead of lymphoid-biased hematopoiesis. Lineage contribution of HSC transduced with PMYs-IRES-EGFP or PMYs-R9-IRES-EGFP retrovirus. E14.5 fetal liver HSPCs (FL-HSPCs) were enriched by Lineage (CD2/3/4/8/B220/CD19/Ter119/Gr1) cell deletion and were infected with PMYs-IRES-EGFP or PMYs-R9-IRES-EGFP retrovirus. Half million HSPC transduced with the related viruses were transplanted into the individual CD45.1 C57BL/6 mice irradiated by X-ray (6.5 Gy). Peripheral blood cells were analyzed 16 weeks after transplantation cells.
